# Biochemical and Structural Characterization of Two Cif-Like Epoxide Hydrolases from *Burkholderia cenocepacia*

**DOI:** 10.1101/2021.01.20.427036

**Authors:** Noor M. Taher, Kelli L. Hvorecny, Cassandra M. Burke, Morgan S.A. Gilman, Gary E. Heussler, Jared Adolf-Bryfogle, Christopher D. Bahl, George A. O’Toole, Dean R. Madden

## Abstract

Epoxide hydrolases catalyze the conversion of epoxides to vicinal diols in a range of cellular processes such as signaling, detoxification, and virulence. These enzymes typically utilize a pair of tyrosine residues to orient the substrate epoxide ring in the active site and stabilize the hydrolysis intermediate. A new subclass of epoxide hydrolases that utilize a histidine in place of one of the tyrosines was established with the discovery of the CFTR Inhibitory Factor (Cif) from *Pseudomonas aeruginosa*. Although the presence of such Cif-like epoxide hydrolases was predicted in other opportunistic pathogens based on sequence analyses, only Cif and its homologue aCif from *Acinetobacter nosocomialis* have been characterized. Here we report the biochemical and structural characteristics of Cfl1 and Cfl2, two <u>**C**</u>i<u>**f**</u>-<u>**l**</u>ike epoxide hydrolases from *Burkholderia cenocepacia*. Cfl1 is able to hydrolyze xenobiotic as well as biological epoxides that might be encountered in the environment or during infection. In contrast, Cfl2 shows very low activity against a diverse set of epoxides. The crystal structures of the two proteins reveal quaternary structures that build on the well-known dimeric assembly of the α/β hydrolase domain, but broaden our understanding of the structural diversity encoded in novel oligomer interfaces. Analysis of the interfaces reveals both similarities and key differences in sequence conservation between the two assemblies, and between the canonical dimer and the novel oligomer interfaces of each assembly. Finally, we discuss the effects of these higher-order assemblies on the intra-monomer flexibility of Cfl1 and Cfl2 and their possible roles in regulating enzymatic activity.

## Introduction

Epoxide hydrolases serve a variety of cellular functions such as detoxification, cell-cell signaling, cell envelope synthesis, and antibiotic resistance. These enzymes act by catalyzing the conversion of a variety of epoxides to their corresponding vicinal diols. Epoxide hydrolases that belong to the α/β hydrolase superfamily of enzymes utilize an Asp-His-Asp/Glu catalytic triad to open the epoxide ring substrate and release the vicinal diol product. In addition to the catalytic triad, the active site of these enzymes usually contains two conserved tyrosine residues (the “Tyr-Tyr pair”) that orient the substrate epoxide ring in the active site pocket and form an oxyanion hole to stabilize a hydrolysis intermediate [1]. However, a new subclass of epoxide hydrolases that utilize a His-Tyr pair instead of the canonical Tyr-Tyr pair was revealed with the discovery of the <u>C</u>FTR <u>I</u>nhibitory <u>F</u>actor (Cif) from *Pseudomonas aeruginosa* [2].

Cif is an epoxide hydrolase from *P. aeruginosa* that serves as a virulence factor in lung infections. Cif has been shown to decrease the concentration of the <u>C</u>ystic <u>F</u>ibrosis <u>T</u>ransmembrane Conductance <u>R</u>egulator (CFTR) on the apical surface of lung epithelial cells, sabotage the inflammation resolution signaling between the epithelial cells and the immune system, and hinder mucociliary clearance of bacteria [3,4,5]. The crystal structure of Cif showed that it utilizes a His-Tyr pair instead of the canonical Tyr-Tyr pair for epoxide ring orientation and hydrolysis intermediate stabilization [6]. <u>a</u>Cif from *<u>A</u>cinetobacter nosocomialis* was subsequently shown to belong to this new class of <u>**C**</u>i<u>**f**</u>-<u>**l**</u>ike (Cfl) epoxide hydrolases [7]. While Cif and aCif have been characterized structurally and biochemically, Cfls present in other opportunistic pathogens remain uncharacterized.

*Burkholderia cenocepacia* is a gram-negative bacterium that belongs to the *Burkholderia cepacia* complex. *B. cenocepacia* encounters diverse environments where its presence has important implications. It is commonly found in the soil where it can cause plant infections, serve as a biocontrol agent to prevent plant infections, and degrade aromatic hydrocarbon compounds [8,9,10,11]. *B. cenocepacia* can also colonize the lung extracellular milieu of immunocompromised and non-immunocompromised patients where it causes high morbidity and mortality due to its broad antibiotic resistance [12,13]. The bacteria are also able to cause a systemic infection by entering the patient’s bloodstream, a condition known “Cepacia Syndrome.” *B. cenocepacia* therefore seems well adapted for survival in a wide range of environments where it must process and respond to a variety of signals and metabolites, such as epoxides. However, no epoxide hydrolases from *B. cenocepacia* have been characterized to date. Additionally, whereas it is known that *P. aeruginosa* can use Cif to hydrolyze 14,15-EET to intercept downstream pro-resolution effects and exacerbate the host’s inflammatory response, it is not known whether *B. cenocepacia* possesses Cfls with such anti-resolution potential.

Here we characterize two predicted Cfls from *B. cenocepacia,* named Cfl1 and Cfl2. We demonstrate that Cfl1 possesses epoxide hydrolysis activity against xenobiotic and potential host-derived epoxides. Our hydrodynamic and structural studies revealed that Cfl1 and Cfl2 exist as octamers and decamers, respectively, unlike Cif and aCif, which exist as dimers. Although the overall assembly of each oligomer is similar, detailed structural analysis revealed key differences between the oligomer as well as the dimer interfaces of each assembly. We finally discuss the possible role of the differential steric constraints placed on the monomer subunit in each structure in regulating enzyme activity.

## Results

### Bcen_3967 and Bcen_4419 are Cif-like predicted epoxide hydrolases

In *B. cenocepacia* strain HI2424, the two proteins with the highest amino-acid sequence identity to Cif are encoded by Bcen_3967 (34% identical, hereafter named Cfl1) and Bcen_4419 (32% identical, hereafter named Cfl2). Amino-acid sequence alignment with Cif shows that both proteins are predicted to have the catalytic residues required for epoxide hydrolysis (Figure 1). Cfl1 and Cfl2 also appear to utilize a Cif-like His-Tyr epoxide ring-opening pair instead of the classical Tyr-Tyr pair. In addition to the catalytic residues, canonical epoxide hydrolases are also distinguished by an H-G-X-P motif, where X tends to be an aromatic residue and P is a *cis*-proline [14]. The H-G-W-P sequence of Cfl1 follows the canonical H-G-X-P motif, whereas Cfl2 substitutes the histidine with an alanine (A-G-F-P). Finally, whereas Cif has an N-terminal secretion signal to direct its export from the cell, neither Cfl1 nor Cfl2 are predicted to have such signal (Figure 1).

**Figure 1.**
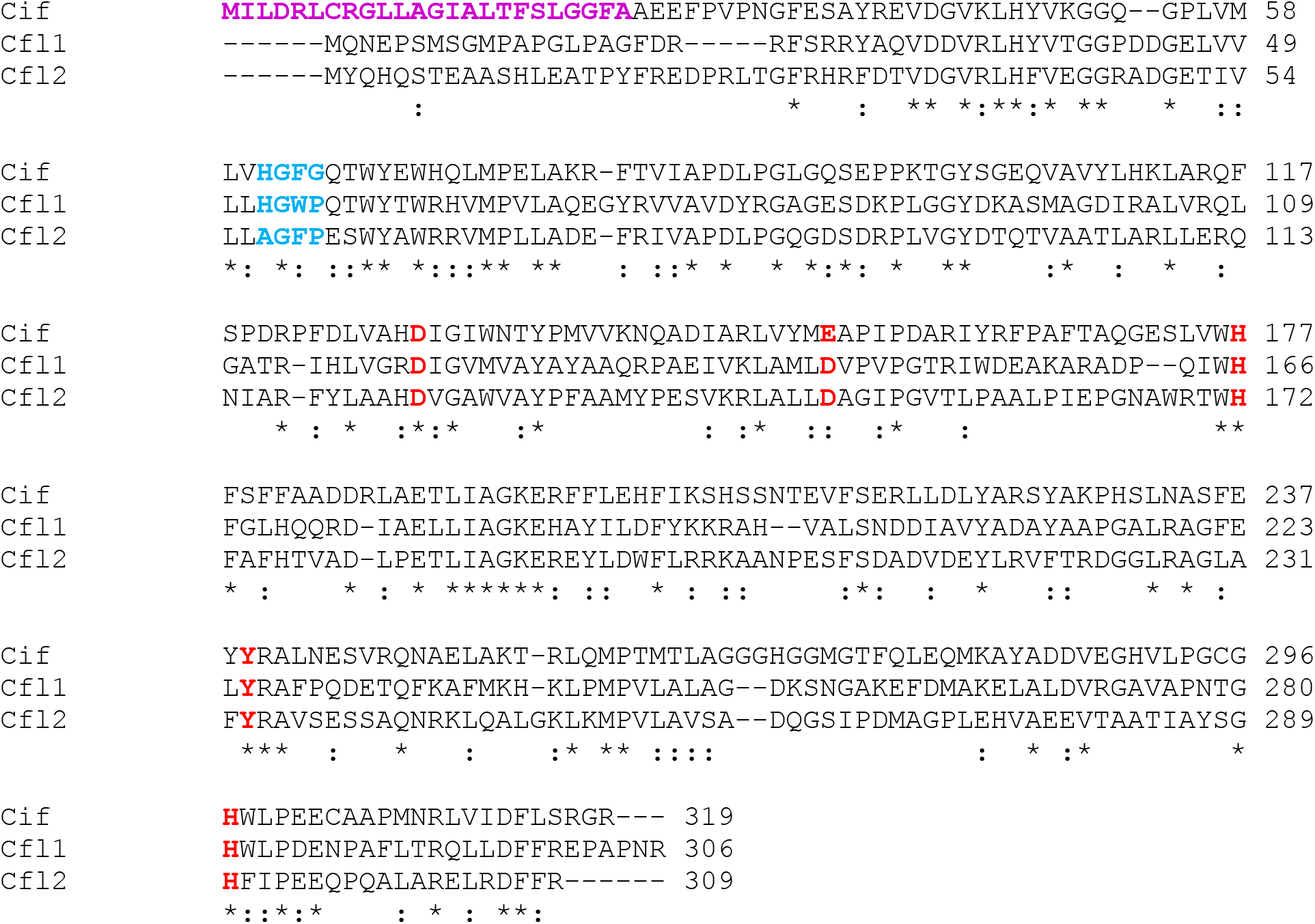
Cfl1 and Cfl2 amino-acid sequences contain Cif-like epoxide hydrolase sequence motifs. ClustalW was used to align the amino-acid sequences of Cif from *P. aeruginosa* strain PA14 with Cfl1 and Cfl2 from *B. cenocepacia* strain HI2424. The residues corresponding to the canonical epoxide hydrolase H-G-X-P motif and the core catalytic residues are highlighted in bold blue and bold red, respectively. The secretion signal sequence of Cif is highlighted in bold magenta. Asterisks and colons below each position indicate identity and similarity among the three sequences, respectively.

### *B. cenocepacia* HI2424 upregulates the transcription of *Cfl2*, but not *Cfl1*, in response to certain epoxides

In *P. aeruginosa* strain PA14, *cif* is part of a three-gene operon that is under regulation by CifR, a TetR-like transcriptional repressor. CifR binds to the intergenic region between *cifR* and the *cif* operon and represses transcription of the operon. The presence of certain epoxides in the media causes CifR to release the DNA and de-repress *cif* transcription [15,16]. *A. nosocomialis* uses a similar epoxide-sensitive regulatory circuit to control the transcription of its *cif* gene [7]. Similarly, *cfl1* and *cfl2* each reside near a gene coding for a TetR-like protein (Figure 2(A)). However, unlike the divergent transcription seen for *cifR* and *cif*, both the *cfl1/cfl1R* and *cfl2/cfl2R* genes appear to be transcribed convergently, and two other members of the *cif* operon in PA14 - *morB* and the MFS transporter gene - are absent from the putative operons of *cfl1/2* (Figure 2(A)). Given the similarities and differences between the *cif* and *cfl1/2* operons, we sought to determine whether the transcription of *cfl1* and *cfl2* is similarly upregulated in response to epoxides. We found that exposure of *B. cenocepacia* HI2424 to epibromohydrin (EBH), *R*-styrene oxide (*R*-SO), and *S*-SO increases transcription of *cfl2* by approximately 5 fold, 15 fold, and 3 fold, respectively, relative to the dimethyl sulfoxide vehicle control (DMSO) as measured by RT-qPCR (Figure 2(B)). In contrast, the same treatments do not result in significantly increased transcription of *cfl1* (Figure 2(B)).

**Figure 2.**
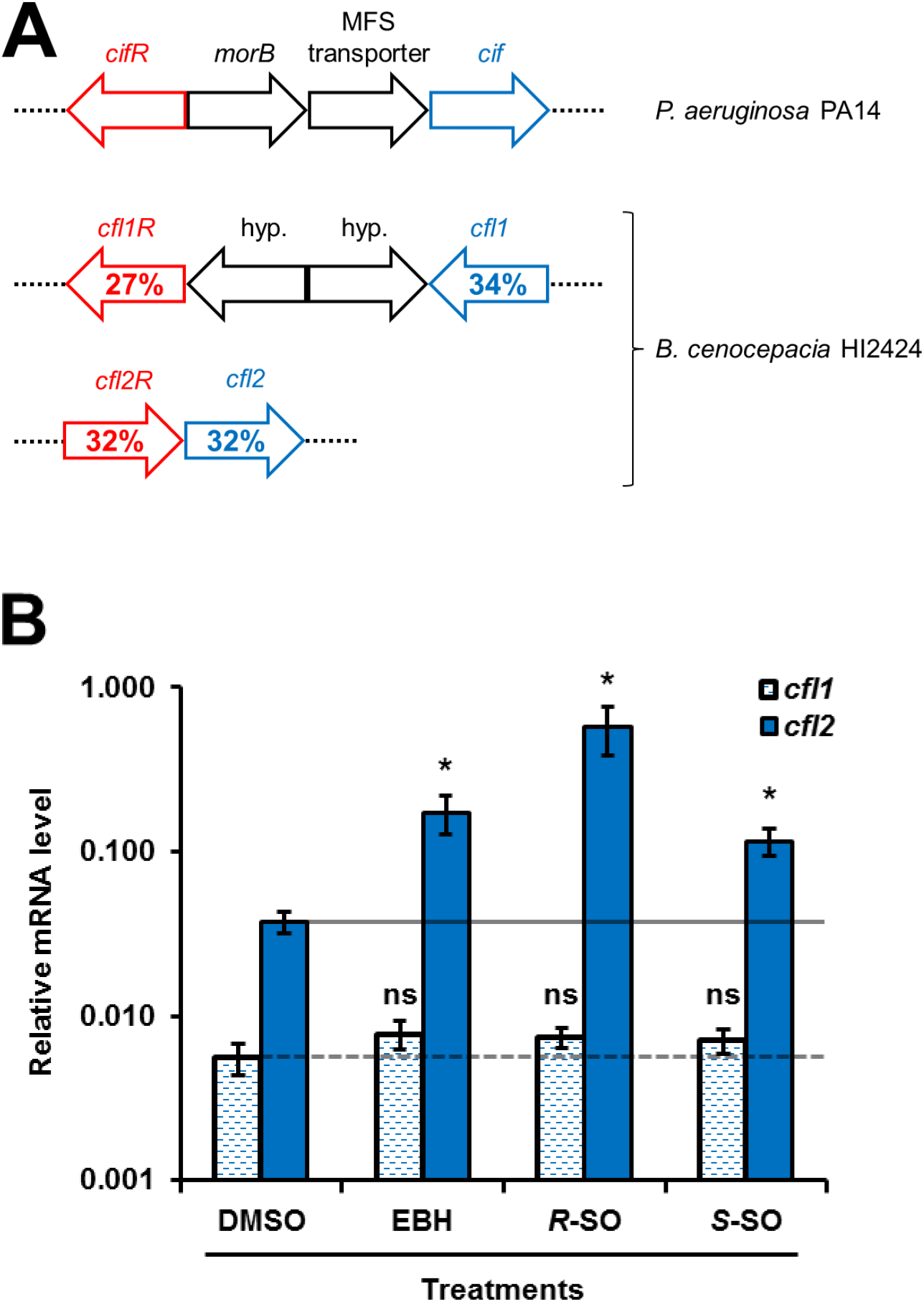
*B. cenocepacia* strain HI2424 upregulates the transcription of *cfl2* in the presence of epoxides in the growth medium. (A) The *cif* operon in *P. aeruginosa* strain PA14 and the genomic regions in *B. cenocepacia* strain HI2424 that encompass *cfl1*-*cfl1R* and *cfl2*-*cfl2R* genes are shown. The percent amino-acid sequence identities of Cfl1 and Cfl2 to Cif (blue) and of Cfl1R and Cfl2R to CifR (red) calculated using EMBOSS Needle [53] are indicated on each respective gene. (B) Relative mRNA levels of *cfl1* and *cfl2* in *B. cenocepacia* strain HI2424 treated with various epoxides (EBH, epibromohydrin; *R*-SO, *R*-styrene oxide; *S*-SO, *S*-styrene oxide) or control (DMSO, dimethylsulfoxide) were measured using RT-qPCR. The change in measured transcript levels of *cfl1* and *cfl2* between treatments was normalized to the levels of *rplU*. Values are reported on a log_10_ scale as the means of three experiments, and the error bars represent the standard deviation. An asterisk indicates *p* < 0.05 when Student’s unpaired *t*-test was used to compare mRNA levels between epoxide-treated and DMSO-treated bacteria (n = 3). ns, not signficant.

### Biochemical characterization of Cfl1 and Cfl2

In order to characterize the enzymatic activities and structures of Cfl1 and Cfl2, we sought to express and purify the two proteins heterologously. Both proteins were overexpressed in *E. coli* BL21(DE3) cells C-terminal to a 10×His-SUMO tag, but only Cfl2 was found in the soluble fraction of the cell lysate. After unsuccessful troubleshooting attempts to obtain soluble Cfl1, we shifted our strategy towards finding a Cfl1 homolog in other *B. cenocepacia* strains with high sequence identity and soluble expression. We found the desired properties in CAR55383.1 from *B. cenocepacia* strain J2315, which exhibits 92% amino-acid sequence identity to Cfl1 (Figure S1). Hereafter “Cfl1” refers to Cfl1 from strain J2315.

Following scarless tag cleavage by the SUMO protease Ulp1 [17], Cfl1 and Cfl2 both eluted from a size exclusion chromatography (SEC) column as large species, with Cfl1 eluting later than Cfl2 (Figure 3(A)). Calibrating the Superdex 200 10/300 SEC column with standards of known relative molar mass (M_r_) revealed that Cfl1 and Cfl2 peak elution volumes fall between those of apoferritin (440 kDa) and β-amylase (200 kDa), with predicted M_r_ values of 240 kDa and 350 kDa, respectively. Since the apparent M_r_ estimates obtained from SEC can be influenced by the shape of the particles in addition to their actual mass, we combined the diffusion coefficients obtained from SEC with the sedimentation coefficients obtained by velocity sedimentation analytical ultracentrifugation in order to gain a more shape-independent estimate of relative molar masses for Cfl1 and Cfl2 (Figure 3(B) and Eq. 1, Materials and Methods). With a sedimentation coefficient of 10.2 S for Cfl1 and 11.2 S for Cfl2, Equation 1 yields M_r_ estimates of 260 kDa and 320 kDa, respectively. Since the predicted M_r_ for a monomer of both proteins is ~34.5 kDa, these results indicate that Cfl1 and Cfl2 exist as higher order oligomers compared to the dimeric Cif and aCif. Finally, we used circular dichroism spectroscopy to examine the thermal stability of the two proteins and determined that Cfl1 and Cfl2 are both stable at physiologically relevant temperatures with melting points of >90 °C and ~70 °C, respectively (Figure 3(C)).

**Figure 3.**
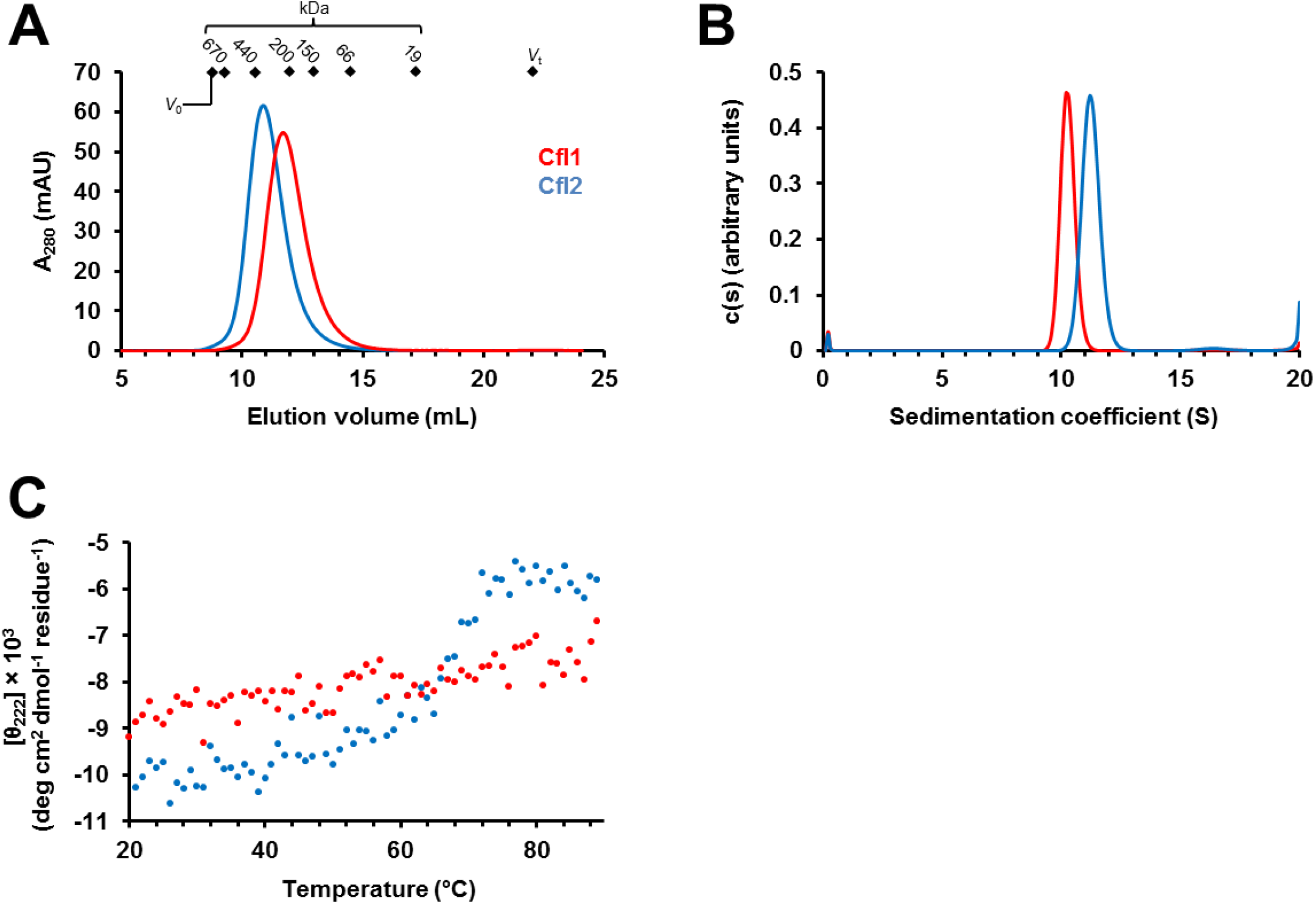
Biochemical characterization of Cfl1 and Cfl2. (A) Cfl1 (red) and Cfl2 (blue) elute as large oligomers in SEC. Diamonds indicate the Superdex S200 10/30 column void volume (*V*_0_), total volume (*V*_t_), and peak elution volumes of molecular mass standards (670 kDa = bovine thyroglobulin, 440 kDa = equine spleen apoferritin, 200 kDa = sweet potato β-amylase, 150 kDa = yeast alcohol dehydrogenase, 66 kDa = bovine serum albumin, 19 kDa = bovine carbonic anhydrase). (B) Sedimentation coefficient distribution of Cfl1 (red) and Cfl2 (blue) analyzed by velocity sedimentation analytical ultracentrifugation. (C) Molar ellipticity at 222 nm (θ_222_) was monitored as Cfl1 (red) and Cfl2 (blue) were heated to determine their melting temperatures (n = 1).

### Epoxide hydrolysis activity of Cfl1 and Cfl2

Given the presence of the epoxide-hydrolase signature motif in the amino-acid sequences of Cfl1 and Cfl2, we performed a preliminary screen for their catalytic activity against a selection of xenobiotic and biologically relevant epoxides. Cfl1 hydrolyzed the *R* enantiomer of SO, and its activity against *S*-SO was significant but nonetheless low compared to the *R*-SO reaction (Figure 4). Cfl1 also showed very low levels of activity against other epoxides (Supplementary Table 1). Surprisingly, Cfl2 had very low activity against all of the tested substrates despite using relatively high protein concentrations (20 to 80 μM) and long incubation periods (1 to 2 hrs) (Figure 4, Table S1). The level of Cfl2 activity against the tested substrates is comparable to the level of activity aCif showed against epoxides that were deemed non-substrates for that epoxide hydrolase [7]. Cfl1 amino-acid sequence alignment with Cif suggests that Asp123 is responsible for the nucleophilic attack that opens the epoxide ring. To test this hypothesis, we expressed and purified a mutant Cfl1 in which Asp123 was substituted with a serine (Cfl1-D123S). Although hydrolysis of *R*-SO by the mutant was still statistically significant compared to its protein-only control, it is sharply decreased compared to the wild-type hydrolysis of *R*-SO (Figure 4).

**Table 1.**
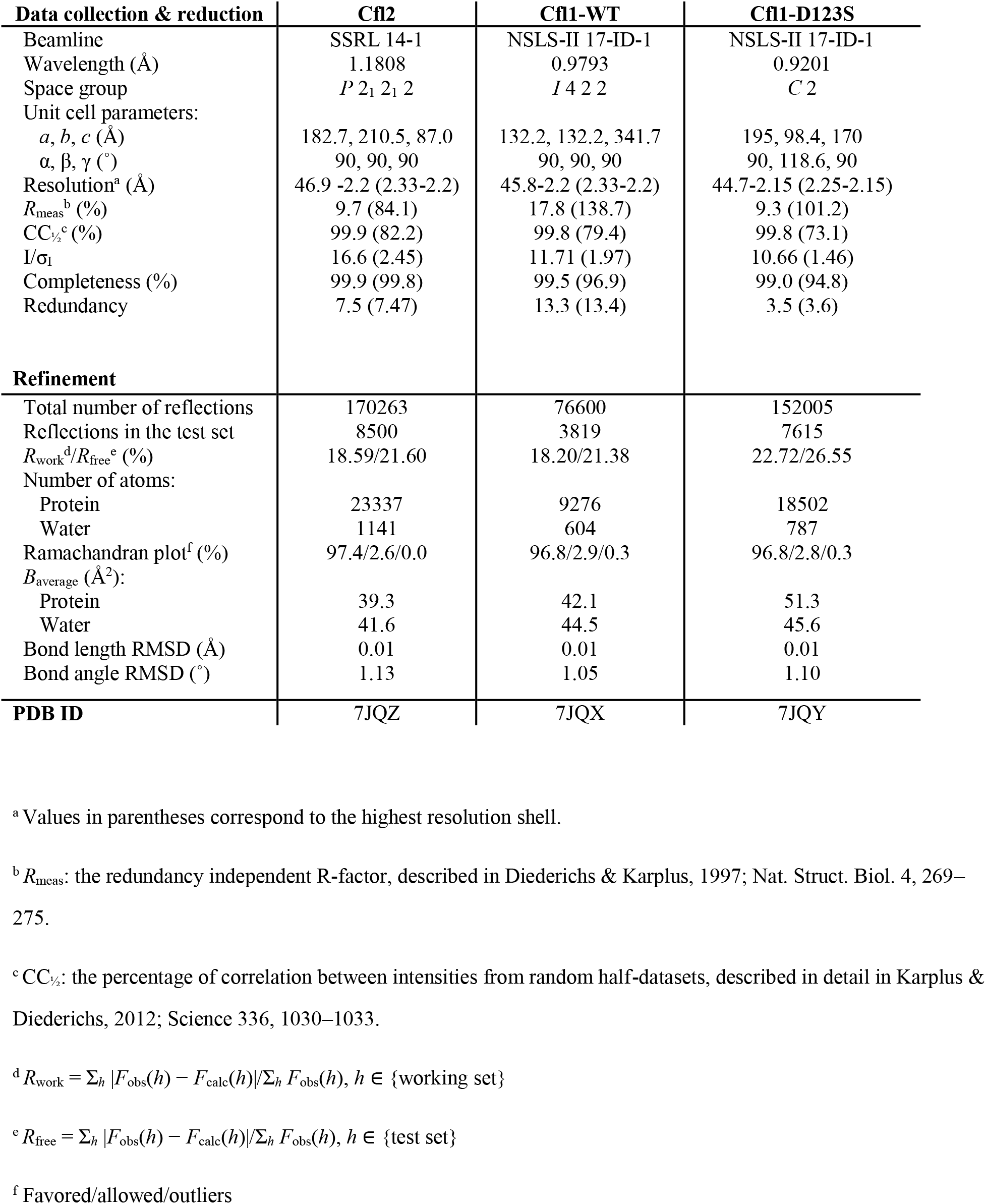
Data collection, reduction, and refinement statistics.

**Figure 4.**
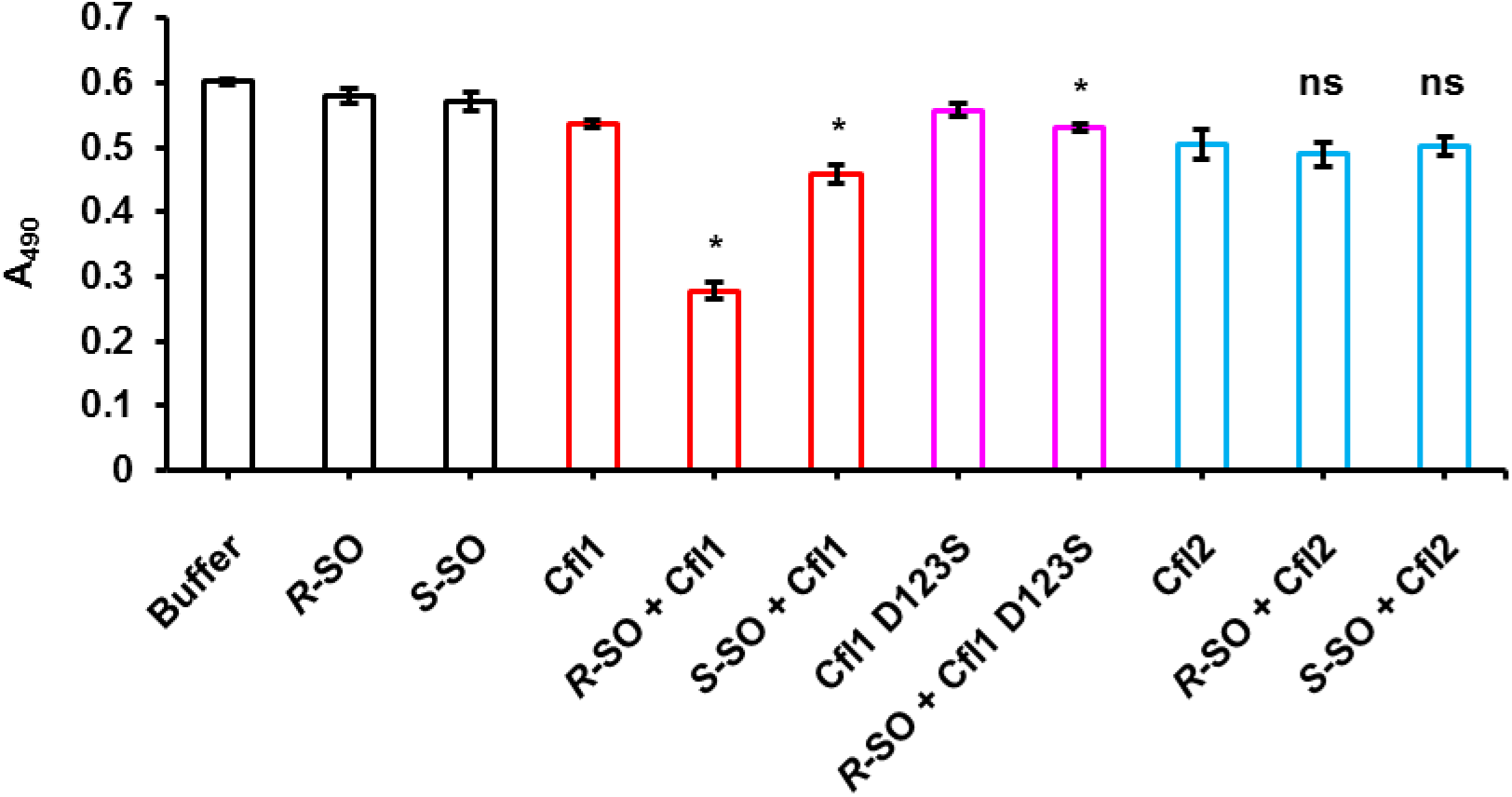
Cfl1 is an active epoxide hydrolase. Cfl1 (red) and Cfl2 (blue) catalytic activity against the *R* and *S* enantiomers of styrene oxide (SO) and the catalytic activity of Cfl1-D123S (magenta) against *R*-SO was determined using the adrenochrome assay. A decrease in absorbance at 490 nm indicates hydrolysis of the epoxide. Values are reported as the means of three experiments, and the error bars represent the standard deviations. An asterisk indicates *p* < 0.05 when Student’s unpaired *t*-test was used to compare the protein+substrate reactions and their respective protein-only controls (n = 3). ns, not signficant.

Cif can convert polyunsaturated epoxide fatty acids upstream of pro-resolving signals into their inactive diol counterparts [18]. *P. aeruginosa* strain PA14 can utilize that capability of Cif to intercept pro-resolving signal pathways between lung epithelial cells and immune cells, ultimately leading to a cycle of host tissue injury and further inflammation [3]. We were interested in determining whether Cfl1 possesses similar activity towards physiologically important host-derived substrates. We compared the hydrolytic activity of wild-type Cfl1 and Cfl1-D123S against a panel of four human-derived epoxides that are known signaling molecules *in vivo* [19,20,21,22]. The four epoxides vary in length, number of double bonds, and the position of the epoxide moiety along the hydrocarbon chain. Wild-type Cfl1 showed significantly more activity against these epoxides compared to Cfl1-D123S (Figure 5).

**Figure 5.**
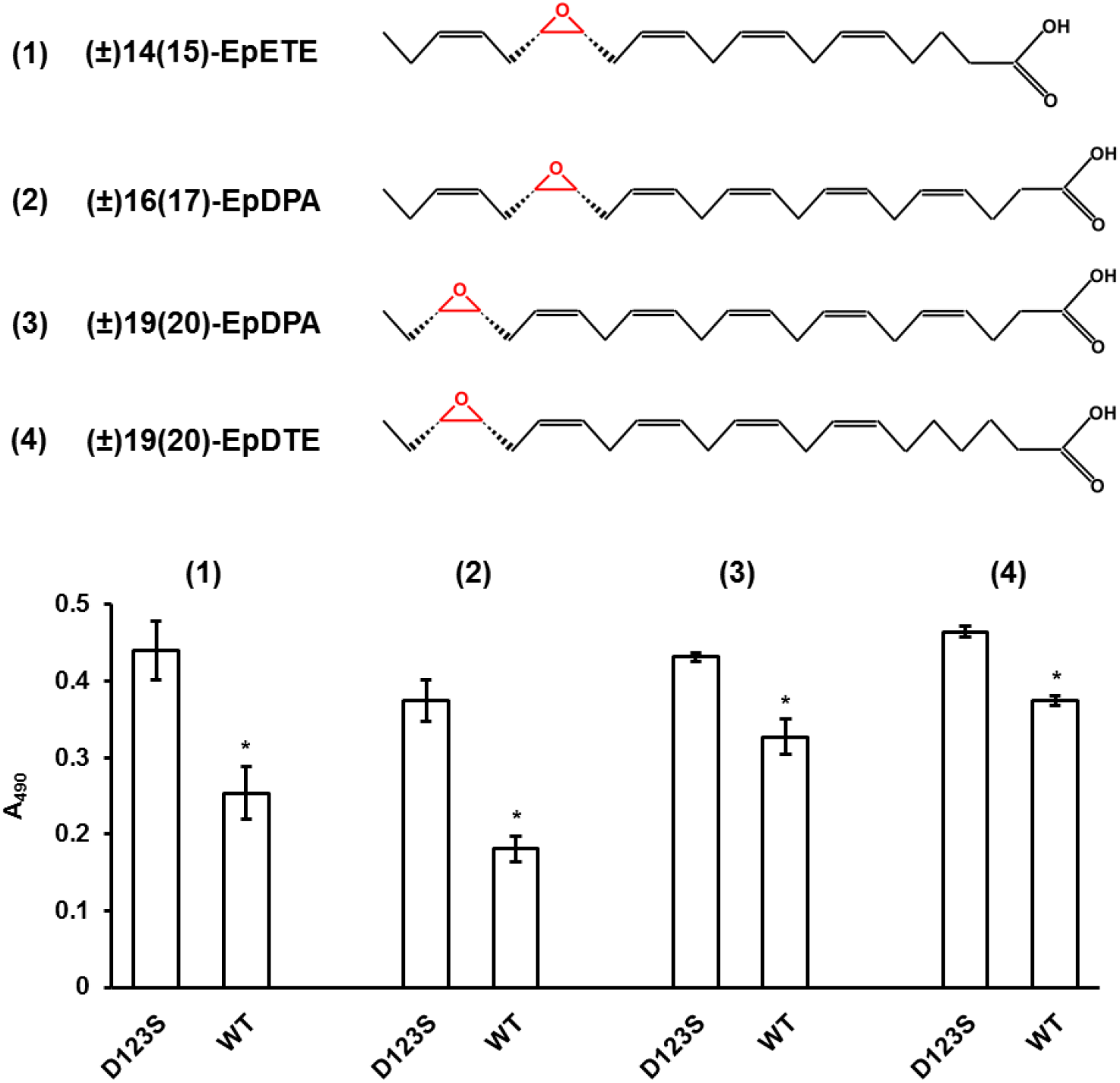
Cfl1 hydrolyzes physiological epoxides *in vitro*. Top: The structures of four physioloical epoxides (1-4) are shown. The epoxide moiety is highlighted in red. Bottom: the hydrolysis activity of Cfl1-D123S and Cfl1-WT against the four physioloical epoxides is shown. Values are reported as the means of three experiments, and the error bars represent the standard deviation. An asterisk indicates *p* < 0.05 when Student’s unpaired *t*-test was used to compare the wild-type enzyme reactions to D123S mutant controls for a particular substrate (n = 3).

### Structural characterization of Cfl1 and Cfl2

As a first look at the architecture of these proteins, we used electron microscopy of negatively stained single particles to examine the overall shape of Cfl2. As shown in Figure S2, a representative raw micrograph of Cfl2 particles shows that they adopt a ring-like structure with a central hole, seen when these molecules adhere to the grid in a “face-on” orientation. Class averages confirm this observation, with some class averages also showing a rectangular shape which is presumably the “side-on” orientation (Figure S2).

In order to obtain a more detailed picture of the structures of Cfl1 and Cfl2, we determined the crystal structures of both proteins to 2.2 Å resolution (Table 1). Each monomer of Cfl1 and Cfl2 exhibits the typical tertiary structure of the α/β hydrolase superfamily, with the main-chain atoms RMSD between Cfl1 and Cif being 0.8 Å and between Cfl2 and Cif being 1.2 Å (Figure 6(A) and Figure 6(B)). When we compared the Cfl1 and Cif active sites, we observed two unique features in Cfl1 (Figure 6(A) inset). First, Asp147, the Cfl1 residue presumed to be the charge-relay acid according to the pairwise sequence alignment with Cif, is not hydrogen bonded with His284, as is typical in acid-base-nucleophile catalytic triads. Instead, that hydrogen bond is satisfied by Ser258. The side chain of Asp147 is oriented in the opposite direction compared to Cif’s Glu153 and forms a salt bridge with Arg122, which is equivalent to His128 in Cif. The second unique feature of the Cfl1 active site is that Arg122 forms a hydrogen bond with the catalytic water. In Cif, the equivalent residue His128 does not form a hydrogen bond with the catalytic water. In contrast to Cfl1, the catalytic side chains of Cfl2 show a much more similar alignment those of Cif (Figure 6(B) inset).

**Figure 6.**
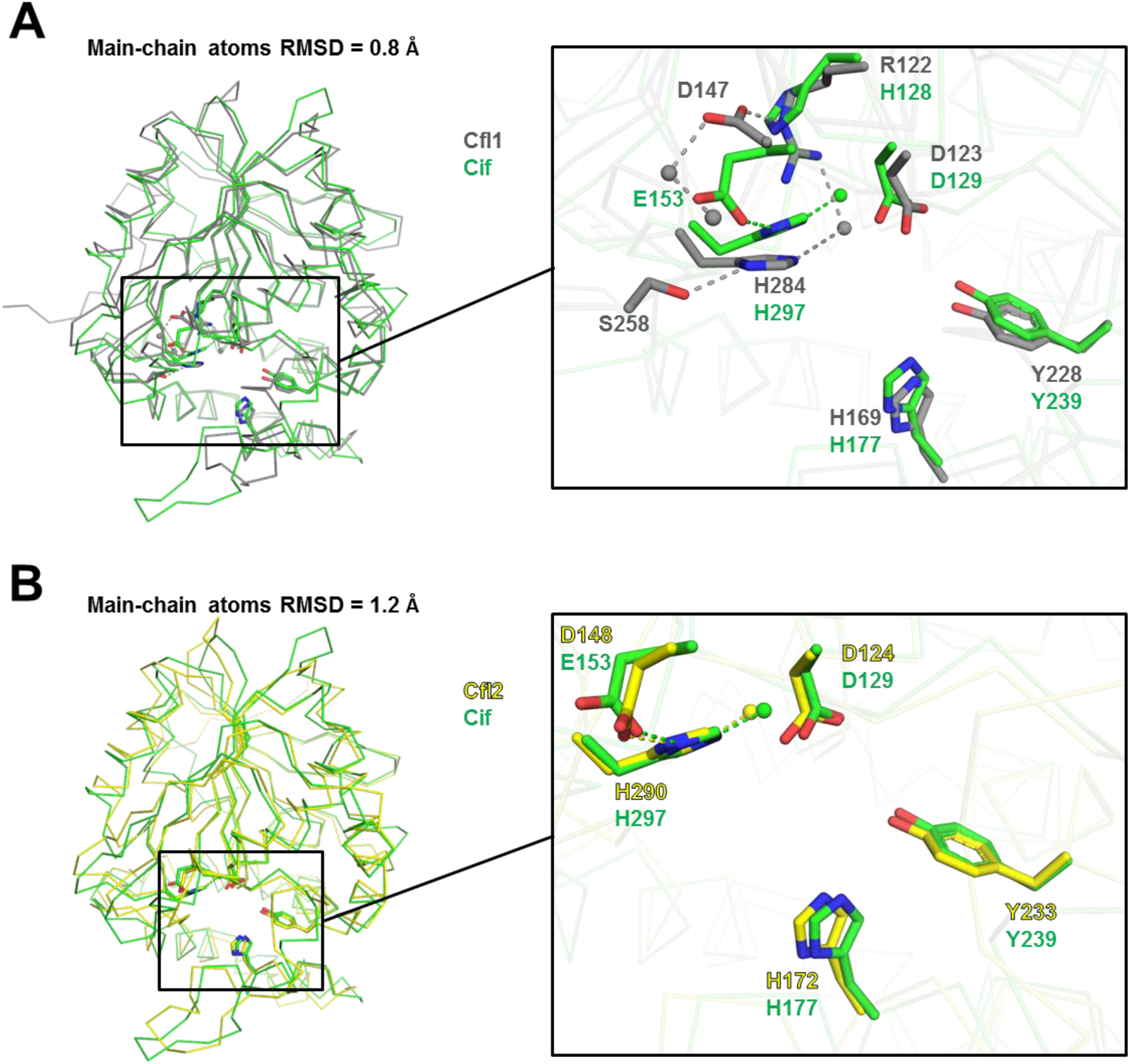
Alignment of the Cfl1 and Cfl2 monomers to the Cif monomer. The main-chain atoms of the Cif (green) and Cfl1 (grey) (A), and of Cif and Cfl2 (yellow) (B) monomers aligned using PyMol are shown. The insets show a closer view of the catalytic and other select active site residues (sticks) and waters (spheres). Electrostatic interactions are represented as dashed lines.

Consistent with the negative-stain electron microscopy and hydrodynamic studies, Cfl1 and Cfl2 form higher order oligomers, either octamers or decamers, respectively, arranged in a ring-like fashion (Figure 7(A) and Figure 7(C)). Each assembly is formed by a ring of dimers; a tetrameric ring in the case of Cfl1, and a pentameric ring in the case of Cfl2. The dimeric subunits of Cfl1 and Cfl2 that are equivalent to the Cif dimer are highlighted in Figures 7(B) (grey) and Figure 7(D) (yellow). Like the Cif dimer, the Cfl1 and Cfl2 dimer subunit is formed through cap domain interactions between two monomers. Unlike Cif, however, each monomer in Cfl1 and Cfl2 has additional oligomerization interfaces that drive the cyclization of the dimer subunits to form the ring-like structures.

**Figure 7.**
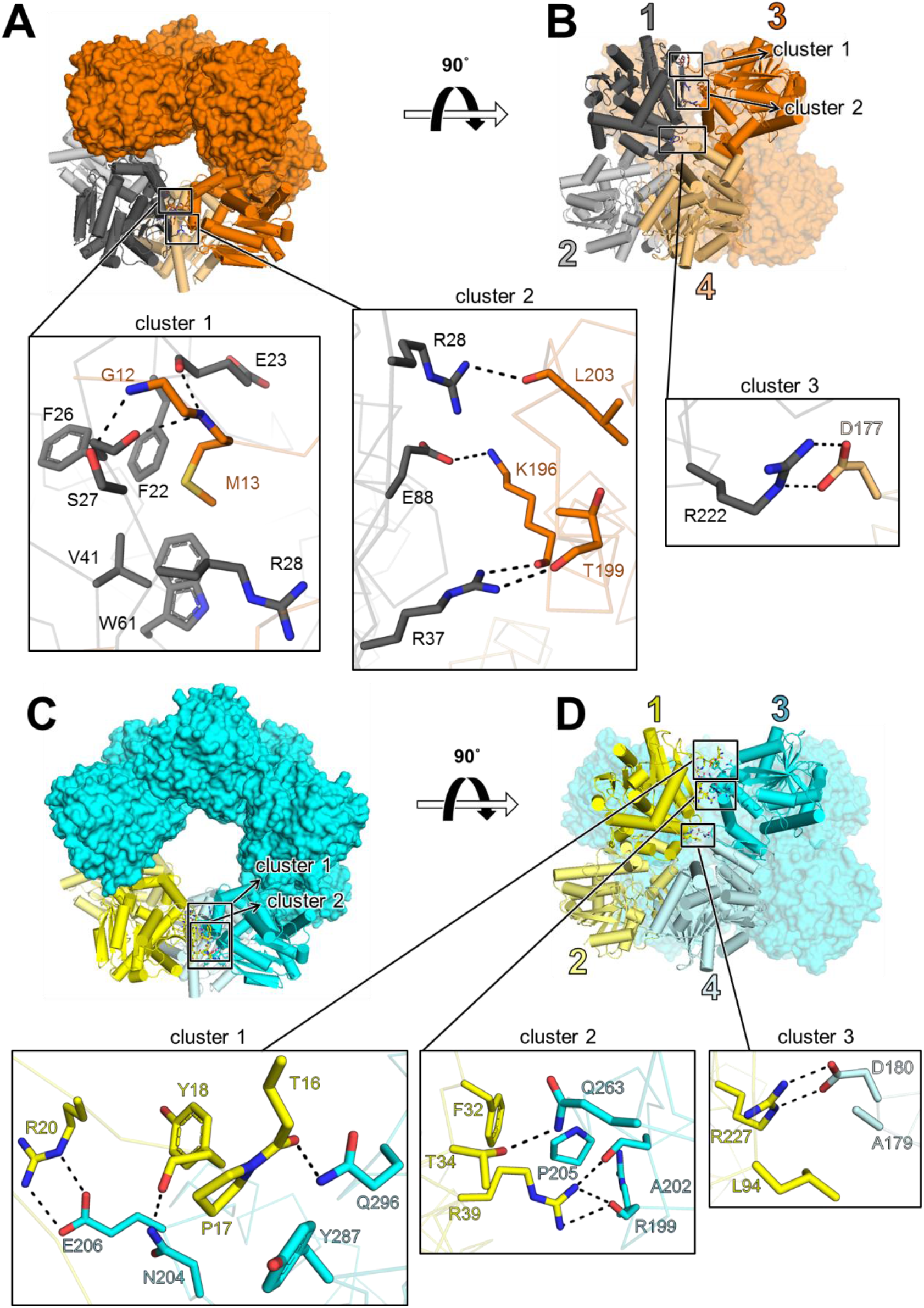
Crystal structures of Cfl1 and Cfl2. (A) Top view of Cfl1. All subunits are shown in surface representation except subunits 1 – 4, which are shown in cartoon representation with cylindrical α-helices. All subunits are colored orange except dimer subunits 1 and 2, which are highlighted in grey. (B) Side view of Cfl1. Colors and representations are the same as in (A). Subunits 1 - 4 are numbered. The positions of Cluster 1 and Cluster 2 in this perspective are only outlined with boxes. (C) Top-down view of Cfl2. All subunits are shown in surface representation except subunits 1 – 4, which are shown in cartoon representation with cylindrical α-helices. All subunits are colored cyan except dimer subunits 1 and 2, which are highlighted in yellow. The positions of Cluster 1 and Cluster 2 in this perspective are outlined with boxes. (D) Side view of Cfl2. Colors and representations are the same as in (C). Subunits 1 - 4 are numbered. Insets represent magnified views of clusters 1 - 3 in each structure. Side chains are depicted as sticks and electrostatic interactions as dashes. Main-chain atoms involved in electrostatic interactions are depicted as sticks; otherwise they are shown as transparent ribbons. Oxygens are colored red, and nitrogens are colored blue.

For an initial, higher-level comparison between the Cfl1 and Cfl2 oligomers, we analyzed their interfaces using Rosetta (Table 2) [23,24,25,26]. The most striking difference is in the dimer (1:2) interface, which is 1080 Å^2^ larger in Cfl2 and has a substantially more favorable binding energy. Given this striking difference, we also analyzed the Cif and aCif dimers. We found that the Cfl2 dimer interface corresponds roughly to those of Cif and aCif, albeit with a modestly higher contribution from hydrophobic residues, whereas the Cfl1 dimer interface is markedly smaller than all of the others (Table 2). In contrast, when comparing the Cfl1 and Cfl2 oligomer interfaces, we found that those of Cfl1 are collectively 435 Å^2^ larger than those of Cfl2 (Table 2).

**Table 2.**
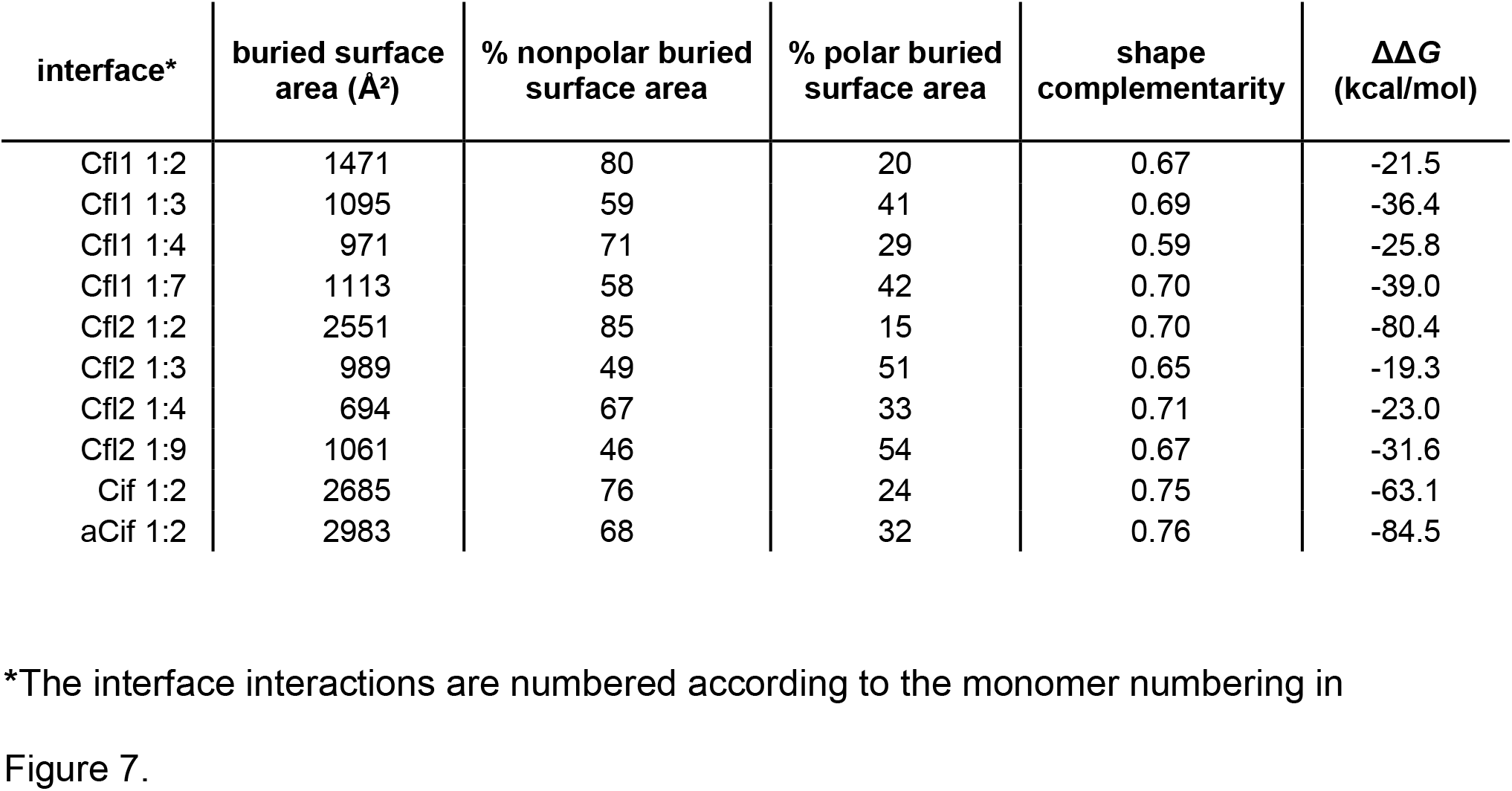
Rosetta analysis of the interfaces of Cfls and Cifs

| interface* | buried surface<br>area (Å<sup>2</sup>) | % nonpolar buried<br>surface area | % polar buried<br>surface area | shape<br>complementarity | ΔΔG<br>(kcal/mol) |
| --- | --- | --- | --- | --- | --- |
| Cfl1 1:2 | 1471 | 80 | 20 | 0.67 | -21.5 |
| Cfl1 1:3 | 1095 | 59 | 41 | 0.69 | -36.4 |
| Cfl1 1:4 | 971 | 71 | 29 | 0.59 | -25.8 |
| Cfl1 1:7 | 1113 | 58 | 42 | 0.70 | -39.0 |
| Cfl2 1:2 | 2551 | 85 | 15 | 0.70 | -80.4 |
| Cfl2 1:3 | 989 | 49 | 51 | 0.65 | -19.3 |
| Cfl2 1:4 | 694 | 67 | 33 | 0.71 | -23.0 |
| Cfl2 1:9 | 1061 | 46 | 54 | 0.67 | -31.6 |
| Cif 1:2 | 2685 | 76 | 24 | 0.75 | -63.1 |
| aCif 1:2 | 2983 | 68 | 32 | 0.76 | -84.5 |

\*The interface interactions are numbered according to the monomer numbering in Figure 7.

We hypothesized that the large difference between Cfl1’s dimerization surface area and energetics and those of Cfl2, Cif, and aCif would allow for more intra-dimer flexibility for Cfl1. In order to test this hypothesis, we carried out Normal Mode Analysis (NMA) to observe the large-scale, longer time-regime movements available for each dimer. In the case of Cfl1 and Cfl2, the dimers were extracted from their respective oligomers before doing the analysis. Unsurprisingly, a comparison of the first non-trivial mode of each dimer shows that the Cfl1 dimer exhibits the largest excursions (Supplementary Movie 1).

When comparing the Cfl1 and Cfl2 structures more closely, we observed that the close-contact (<3.2 Å) interactions at their oligomerization interfaces are organized in three similar “clusters” (Figure 7, insets). Clusters 1 and 2 are both part of the in-ring interfaces [*i.e.*, 1:3, 1:7 (Cfl1), and 1:9(Cfl2)], while cluster 3 is part of the cross-ring (*i.e.*, 1:4) diagonal interface. Cluster 1 is where the N-terminus of one monomer contacts the neighboring monomer. In the case of Cfl1, monomer 1 interacts with the N-terminus of adjacent monomer 3, but in Cfl2 the opposite is true: monomer 3 interacts with the N-terminus of monomer 1 (Figure 7(A) and Figure 7(D), cluster 1; Figure S3(A) and Figure S3(B)). The different oligomerization states of the two structures coincide not only with different N-terminal donor-acceptor relationships among neighboring monomers within each ring, but also different conformations of the monomers in the dimer subunit across each ring (Figure S3(C) and Figure S3(D)). The N-terminus of Cfl1 monomer 3 makes several electrostatic interactions with monomer 1, as well as a hydrophobic interaction mediated by the insertion of Met13 into a pocket comprised of Phe22, Phe26, Val41, Trp61, and the hydrocarbon portions of Glu23 and Arg28 side chains (Figure 7(A), cluster 1). Similarly, in Cfl2, the N-terminus of monomer 1 makes several electrostatic interactions with monomer 3 and a hydrophobic interaction between Pro17 and Tyr287 (Figure 7(D), cluster 1). Cluster 2 is located near Cluster 1 in both structures but does not involve the N-terminus (Figure 7(A) and Figure 7(D), cluster 2). Lastly, Cluster 3 is where monomer 1 contacts monomer 4, and is the only point of contact between these two monomers in both structures (Figure 7(B) and Figure 7(D), cluster 3).

Finally, we sought to confirm that the sharp decrease in the activity of Cfl1-D123S seen in Figure 4 is not due to a gross change in the overall structure of the enzyme or its active site caused by the single amino-acid substitution. To that end, we determined the crystal structure of Cfl1-D123S to 2.15 Å resolution (Table 1). As shown in Figure S4, the RMSD of main-chain atoms between wild-type Cfl1 and Cfl1-D123S monomers is 0.2 Å. Closer inspection of the mutant enzyme’s active site shows that, apart from the D123S substitution, the active site is largely unchanged compared to the wild-type enzyme (Figure S4, inset).

### Evolutionary conservation of the Cfl1 and Cfl2 interfaces

Given the novel oligomerization interfaces of Cfl1 and Cfl2 in comparison to other structurally characterized members of the α/β hydrolase superfamily, we were interested in examining the evolutionary conservation of the oligomer interfaces and how they compare to the canonical dimer interfaces. We used the ConSurf server to analyze the sequence conservation of Cfl1 and Cfl2 [27,28].

Comparing the average conservation scores of the interfaces’ surface residues reveals that the Cfl1 dimer interface is more conserved than its oligomeric interfaces, while the Cfl2 dimer interface, as a whole, does not appear to be the most conserved interface (Table S2). Both dimer interfaces are more conserved on average than the non-interface surface residues of their respective proteins (Table S2). We note that these differences in the means are not statistically significant due to relatively large standard deviations of conservation scores across whole interfaces. Indeed, comparing the local conservation pattern between the interfaces reveals interesting differences between the dimer interfaces and the oligomer interfaces of both proteins (Figure 8). Both Cfl1 and Cfl2 1:2 interfaces contain a central, relatively larger area of highly conserved residues that is lacking in their oligomeric interfaces. When we analyzed the equivalent 1:2 interfaces of Cif and aCif, we found a similarly positioned and conserved “pivot point” in both dimer interfaces (Figure 8). This suggests that a portion of the dimer interfaces may have evolved before the oligomer interfaces, and that the dimer interface may be under a different evolutionary pressure to serve a different function than the oligomer interfaces. Compared to the Cfl1 and Cfl2 1:2 interface, the Cfl1 1:7 interface and Cfl2 1:9 interfaces, which is where we observe the differential N-terminus donor-acceptor relationship between Cfl1 and Cfl2 monomers, appear to be the least conserved (Figure 8 and Table S2).

**Figure 8.**
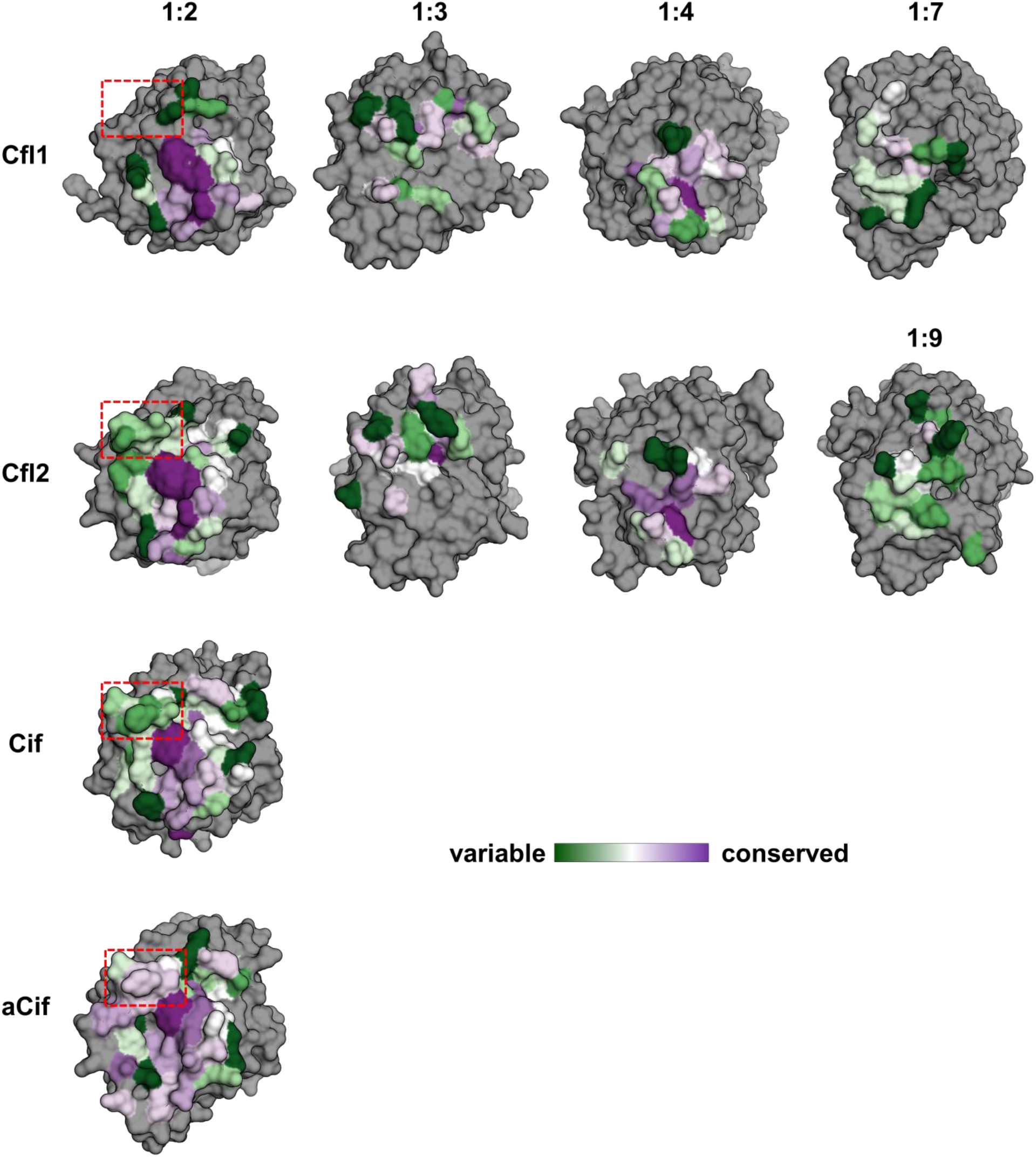
Evolutionary conservation of the interfaces of the Cfls and Cifs. The interfaces of Cfl1, Cfl2, Cif, and aCif are shown on each protein’s monomer subunit as surfaces. The interfaces are labeled at the top according to the monomer numbering used in Figure 7. The interface residues calculated by Rosetta are colored according to the ConSurf conservation score as indicated by the color legend, and all other residues are colored grey. The region outlined with red dashes indicates a structural element that is found in Cfl2, Cif, aCif, but not Cfl1.

Although the dimer interfaces of all four proteins feature a central pivot point that is highly conserved, this region is surrounded by more polymorphic residues in all cases (Figure 8). One such polymorphic patch found Cfl1 and Cfl2 is missing from Cfl1’s interface (Figure 8 red dashes). This missing region in Cfl1 has a dominant contribution to its smaller dimer interface surface area compared to Cfl2, Cif, and aCif. The same area in aCif shows above-average conservation, but remains less conserved than the core of the interface (Figure 8). Along with NMA results showing that the isolated Cfl1 dimer is the most flexible of the four, these data suggest that the dimer interfaces retain a central conserved pivot point with the surrounding polymorphic region serving as a modifier of inter-dimer flexibility according to functional needs.

## Discussion

Structural flexibility is a critical factor in enzymatic catalysis. Flexibility plays an important role in substrate specificity and turnover, and it is perhaps especially important for enzymes in which two or more domains contribute to the active site architecture [29,30]. Indeed, protein scientists have learned from nature to leverage this principle and manipulate enzyme dynamics to create new functionalities [31].

Despite some similarities between the two structures, the steric environments of the Cfl1 and Cfl2 monomers are distinct and likely lead to differences in freedom of motion. The key differences we noted include the N-terminus donor-acceptor relationships between the monomers, the surface areas and binding energies of the interfaces, and the conformations of the dimer subunit in each structure. Comparative NMA between the dimers of Cfl1, Cfl2, Cif, aCif showed that the *ex circulum* (*i.e.*, out of the ring) dimer of Cfl1 with its smallest dimer interface is the most flexible (Supplemental Movie 1). However, within the ring, several factors may act together to further differentiate the steric environment of the Cfl1 monomer from that of Cfl2, and the difference in the dimer interface is but one of them.

Further analysis of the first three non-trivial normal modes of the four proteins reveals that the Cif and aCif dimer motions are coupled to extensive intra-monomer deformations, specifically between the core and cap domains (Figure 9). Compared to Cif and aCif, the extracted Cfl1 dimer deformations are concentrated in a much smaller footprint (Figure 9, “*ex circulum*”). While the dimer interface is smaller in Cfl1, the interfaces formed during its further oligomerization are actually somewhat more extensive than the interfaces formed by Cfl2 (delta = +435 Å^2^), but not to an extent that compensates fully (Table 2). As a result, within the Cfl1 ring, the three dominant modes include interdimer flexion, seen in the form of new areas of deformation which correspond to its oligomer interfaces that are lacking from the *ex circulum* Cfl1 dimer (Figure 9, bottom row). This is not the case with Cfl2, where the *ex circulum* dimer loses its deformability at the dimer interface once it oligomerizes but the oligomer interface deformations do not compensate to the same degree as they do in Cfl1 (Figure 9, bottom row). Correspondingly, within the ring, Cfl1 retains some flexibility at the dimer interface, whereas Cfl2 appears more tightly constrained.

**Figure 9.**
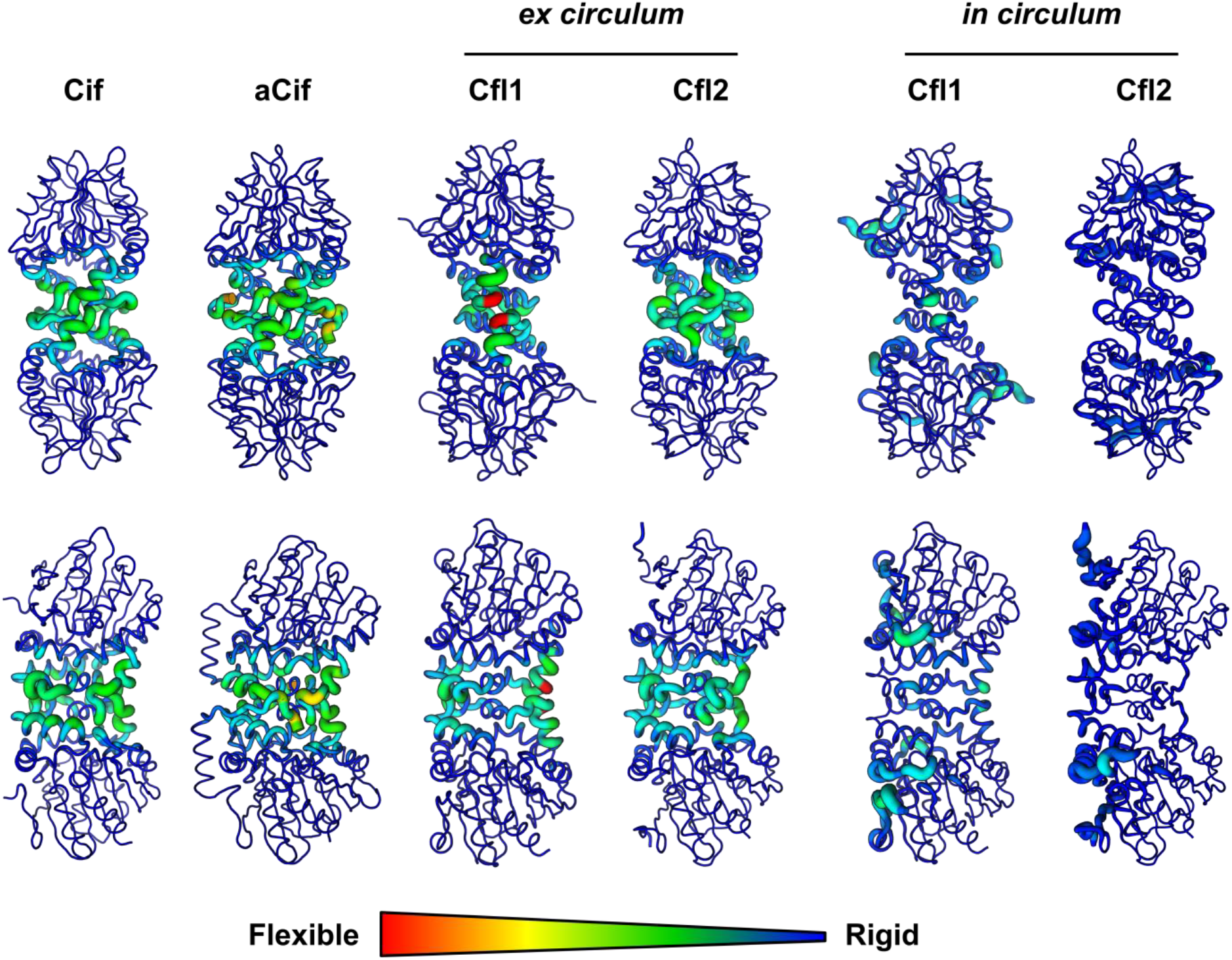
Deformation energy analysis of the Cfls and Cifs. The sum of the deformation energies of the first three modes of each protein is depicted as a heatmap onto its structure. The colors and thickness of the heatmap indicate relative flexibility and rigidity as shown by the scale. The top row shows the face-on view of each dimer, and the bottom row shows the side-on view. In the case of the “*ex circulum*” (out of the ring) Cfl1 and Cfl2 dimers, the calculations were performed on the dimer after extracting it from its respective oligomer. *in circulum*, in the ring.

These observations suggest that the Cfl1 and Cfl2 rings offer different degrees of flexibility for their respective dimers. We hypothesize that in the ring state, Cfl1, but not Cfl2, retains sufficient intra-monomer flexibility to achieve activity. Additionally, we speculate that dissociation or loosening of the ring may modulate the activity of each protein.

The involvement of the N-termini in forming the oligomer interfaces of Cfl1 and Cfl2 represents an extension of the known functions of the N-termini in α/β epoxide hydrolases. In aCif, the N-terminus forms an α-helix on the side of the protein opposite the active site, and then contacts the active-site entry by wrapping around the dimer interface as an unstructured loop, forming monomer-monomer interactions along the way [7]. Interestingly, the position of the aCif monomer-monomer contacts mediated by the N-termini resembles that of the 1:4 (Cluster 3) interface contact in Cfl1 and Cfl2. *Aspergillus niger* epoxide hydrolase has a long N-terminus extension which forms a “meander” that also participates in the dimer-interface contacts [32]. These similarities suggest a possibly more general role for the N-terminus in stabilizing monomer-monomer interactions in some α/β epoxide hydrolases.

*B. cenocepacia* can establish a presence in a diversity of microenvironments, ranging from the soil to the human lung, where the epoxide hydrolysis of Cfl1 may be deployed. Cfl1 may detoxify xenobiotic epoxides encountered by *B. cenocepacia* in these environments, or funnel them as an energy source in a manner similar to other microorganisms [33,34]. In addition to xenobiotic epoxides, we found that Cfl1 can hydrolyze human-derived epoxides. This raises the possibility that *B. cenocepacia* may potentially utilize Cfl1 in a manner similar to *P. aeruginosa*’s use of Cif to sabotage host signaling pathways. Although Cif’s ability to be secreted presumably aids in its capacity to encounter and hydrolyze host epoxides during an infection, it is unknown if secretion is a requirement, as fatty acids are able to passively diffuse through the membrane [35]. Cfl1 is predicted to lack general secretory pathway (Sec) and twin-arginine translocation (TAT) motifs. However, there have been instances in the literature of proteins lacking a strict TAT motif yet are secreted through this pathway directly or indirectly [36,37,38].

The lack of substantial Cfl2 activity raises the possibility of it functioning as a non-catalytic pseudoenzyme *in vivo*. Pseudoenzymes are typically classified as such based on sequence analyses that reveal missing critical catalytic residues. However, this is not the case with Cfl2 which maintains the epoxide hydrolase motifs in its amino acid sequence and 3-dimensional arrangement of its catalytic residues. A previous study by van Loo *et al.* found that four out of twelve predicted bacterial epoxide hydrolases they solubly expressed showed very low hydrolysis activity against the 25-substrate panel they tested [14]. As the number of biochemical studies of epoxide hydrolases continues to accumulate in the future, sequence analyses paired with structural comparisons may allow us to fine tune the criteria for differentiating between active and pseudo-epoxide hydrolases.

## Materials and Methods

### RT-qPCR

*cenocepacia* strain HI2424 was streaked on an lysogeny broth (LB; [39]) agar plate and incubated overnight at 37 °C. Several colonies were used to inoculate a 5 mL LB broth and grown overnight at 37 °C in a rotary incubator. Fifty microliters of the overnight culture was used to inoculate 5 mL of LB broth containing 0.02% DMSO [(*v/v*), =2.8 mM], 1 mM of EBH, *R*-SO, or *S*-SO. When each culture reached an O.D._600_ of 1.0, 1 mL of culture was spun down at 3000 × *g*, 4 °C, for 10 mins and the pellet was stored at −80 °C. RNA was extracted from thawed pellets using the RNeasy RNA extraction kit (Qiagen), and genomic DNA was digested using the RQ1 kit (Promega). cDNA was generated using qScript^®^ cDNA SuperMix (Quantabio) and stored at −20 °C. RT-qPCR was carried out using PerfeCTa^®^ SYBR^®^ Green FastMix^®^ for iQ™. “No-template” and “extracted RNA template” RT-qPCR reactions were included as negative controls for genomic DNA contamination. Genomic DNA from *B. cenocepacia* strain HI2424 was used to generate a standard curve for each pair of primers.

### Cloning, protein expression, and purification

Cfl1 (accession number ABK11134.1) and Cfl2 (accession number ABK10685.1) from *B. cenocepacia* strain HI2424 and Cfl1 from *B. cenocepacia* strain K56-2 (EPZ88246.1, 100% amino acid identity to CAR55383.1 from strain J2315) coding sequences were PCR amplified and inserted into pCDB24 digested with Xho1 using Gibson Assembly. BL21(DE3) cells were transformed with plasmid and grown on LB + 100 μg/mL carbenicillin agar plates at 37 °C overnight. One transformed colony was used to inoculate 10 mL of LB + 100 μg/mL carbenicillin broth and grown overnight at 37 °C. After ~12 hrs, the overnight culture was used to inoculate 1 L of Terrific Broth [40] in a 2 L baffled flask and allowed to grow at 37 °C with shaking at 180 rpm. When the culture reached an O.D._600_ of ~0.3, the bacteria were moved to 16 °C with shaking at 180 rpm for 1 hr. Expression was then induced with 0.1 mM isopropyl β-D-1-thiogalactopyranoside. After ~24 hrs of expression, the bacteria were spun down at 5250 × *g* for 30 mins at 4 °C. The supernatant was discarded and the pellet was resuspended in lysis buffer (500 mM NaCl, 20 mM Tris, pH 8.5, 40 mM imidazole, pH 8.5, 2 mM MgCl_2_) supplemented with 25 units/mL of Pierce Universal Nuclease. Cells lysis was carried out using an M-110L microfluidizer (Microfluidics) in 3 passes at ~18 kpsi. The cell lysate was spun down at 40,000 rpm in a Type 45 Ti rotor for 1 hr at 4 °C, and the supernatant was filtered through a 0.45 μm MCE membrane (Millipore) to remove residual cell debris. Five milliliters of HisPur Ni-NTA resin (Thermo Fisher Scientific) washed with 50 mL of equilibration buffer (500 mM NaCl, 20 mM Tris, pH 8.5, 40 mM imidazole, pH 8.5) was added to the clarified cell lysate, and the mixture was gently stirred at room temperature for 30 min. The mixture of resin and clarified lysate was then passed through a gravity column. The protein-bound resin was washed with 50 mL of equilibration buffer followed by two 50 mL washes with 500 mM NaCl, 20 mM Tris, pH 8.5, 100 mM imidazole, pH 8.5. The protein was then eluted in 50 mL of 500 mM NaCl, 20 mM Tris, pH 8.5, 500 mM imidazole, pH 8.5. The eluate was supplemented with 5.6 mL of 10× Ulp1 reaction buffer (1.5 M NaCl, 500 mM Tris, pH 8.5, 10 mM DTT, 2% [v/v] IGEPAL CA-630) and 1.2 mg of Ulp1 protease. The cleavage reaction was simultaneously dialyzed overnight against 4 L of 20 mM NaCl, 20 mM Tris, pH 8.5, 0.5 mM TCEP-HCl at room temperature. The cleaved protein was then passed through HisPur Ni-NTA resin equilibrated with 20 mM NaCl, 20 mM Tris, pH 8.5, 5 mM imidazole, pH 8.5. The flow-through was then applied to a HiTrap Q HP 5 mL anion exchange chromatography column (GE Healthcare) attached to an ÄKTA Explorer 100 system (Amersham Pharmacia Biotech). The protein was eluted using a 200 mL gradient of wash buffer (20 mM NaCl, 20 mM Tris, pH 8.5) and increasing concentration of elution buffer (1 M NaCl, 20 mM Tris, pH 8.5) [17].

### Hydrodynamic analysis

A Superdex 200 10/300 SEC column (Amersham Pharmacia Biotech) attached to an ÄKTA FPLC system (Amersham Pharmacia Biotech) was used to obtain peak elution volumes for Cfl1, Cfl2, and SEC standards (Sigma Aldrich). Column equilibration and sample elution were carried out at 4 °C using 150 mM NaCl, 50 mM sodium phosphate pH 7.4, and 0.02% (*w/v*) NaN_3_ at a flow-rate of 0.5 mL/min. The diffusion coefficients (D) of Cfl1 and Cfl2 were extrapolated from the plot of the partitioning coefficients (*K*_av_) of the standards versus their known diffusion coefficients.

Velocity sedimentation analytical ultracentrifugation was carried out using a ProteomeLab XL-A centrifuge (Beckman Coulter) at 25,000 rpm, 20 °C, in an An-60 Ti rotor. Sample sedimentation was monitored through absorbance at 280 nm. Data were analyzed using SEDFIT (15.01b) and SEDNTERP (20120828 BETA) to obtain the sedimentation coefficients [41,42]. The relative molar masses of Cfl1 and Cfl2 were estimated using the Svedberg equation:

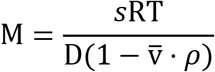

where M is the relative molar mass of the protein in g/mol, *s* is the sedimentation coefficient of the protein in Svedbergs (10^−13^ sec), R is the gas constant in J/mol/K, T is the temperature in K, D is the diffusion coefficient of the protein in cm^2^/sec, 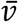 is the partial specific volume of the protein in mL/g, and *ρ* is the buffer density in g/mL.

### Circular dichroism spectroscopy

Circular dichroism spectroscopy was performed using a JASCO J-815 spectrometer equipped with a CDF-426S Peltier temperature controller and a 5 mm-pathlength quartz cuvette. Circular dichroism spectroscopy was monitored at 222 nm for a buffer-only sample (20 mM sodium phosphate, pH 7.4) and 1.5 μM Cfl1 or Cfl2 samples while the temperature was increased from 20 to 90 °C in 1 °C increments with a 5-second period of equilibration before taking each measurement. The raw data of the buffer-only sample were subtracted from the raw data of the protein samples before plotting the molar ellipticity.

### Epoxide hydrolysis assay

An adrenochrome reporter end-point assay adapted from Cedrone *et al.* was used to measure the relative amount of hydrolyzed epoxide [43]. Absorbance at 490 nm was measured using a TECAN Infinite M1000 reader or a Biotek Synergy Neo2 reader. All reactions were carried out at 37 °C with 20 μM protein and 2 mM substrate in 100 mM NaCl, 20 mM HEPES, pH 7.4, and 2% DMSO. When testing the activity of Cfl2 against biological substrates, 80 μM protein was used. The xenobiotic epoxide reactions proceeded for 1 hr in a 100 μL reaction, and the poly-unsaturated fatty acid epoxide reactions proceeded for 2 hrs in a 50 μL reaction. The reactions were quenched by adding 0.5× the starting reaction volume of a solution of 90% acetonitrile containing sodium periodate at a 1:1 molar ratio to the starting substrate and incubating for 30 mins at room temperature. A molar excess of epinephrine-HCl in 0.5× the starting reaction volume was finally added, and an aliquot of 100 μL (xenobiotic epoxide reactions) or 90 μL (fatty acid epoxide reactions) was then used to measure absorbance at 490 nm in a transparent 96-well flat-bottom plate.

### Electron microscopy

Using a 400-mesh carbon-coated (4-6 μm thick) copper grid that was glow discharged for 30 secs at 20 mA in residual air, a five microliter aliquot of Cfl2 at a concentration of 10 μg/mL in 100 mM NaCl, 20 mM HEPES, pH 7.4 was applied to the grid for 30 seconds. The solution was then quickly blotted with filter paper, briefly washed with 5 μL of buffer and blotted, and then stained with 5 μL of 0.75% (*w/v*) uranyl formate, blotted, and then left to air dry. Images were acquired on an FEI Tecnai F20ST microscope with a Gatan OneView camera. Reference-free class averages of 800 manually picked particles were generated using EMAN2 [44].

### Crystallography

Cfl2 crystals were obtained in a hanging drop composed of 2 μL of 1.5 mg/mL protein in 20 mM NaCl, 10 mM HEPES pH 7.5 and 2 μL of well solution (784 mM sodium thiocyanate, 12% PEG 3350) and equilibrated by vapor diffusion against 400 μL of well solution. The harvested crystals were soaked in cryoprotectant solution (20% ethylene glycol, 800 mM sodium thiocyanate, 14% PEG 3350) before flash cooling in liquid nitrogen. The oscillation data collection was performed at the Stanford Synchrotron Radiation Lightsource beamline 14-1 equipped with a MAR325 detector at 100 K, a phi of 0.3°, a total of wedge of 180°, and a wavelength of 1.18076 Å. Wild-type Cfl1 crystals were obtained in a hanging drop composed of 2 μL of 5.2 mg/mL protein in 20 mM NaCl, 20 mM HEPES pH 7.4 and 2 μL of well solution (750 mM potassium sodium tartrate, 100 mM HEPES pH 8.1) and equilibrated by vapor diffusion against 400 μL of well solution. The harvested crystals were directly flash cooled in liquid nitrogen without further cryoprotection. The oscillation data collection was performed at the National Synchrotron Light Source II beamline 17-ID-1 (AMX) equipped with an Eiger 9M detector at 100 K, a phi of 0.1°, a total of wedge of 180°, and a wavelength of 0.979331 Å. Cfl1-D123S crystals were obtained in a hanging drop composed of 2 μL of 1.9 mg/mL protein in 20 mM NaCl, 20 mM HEPES pH 7.4 and 2 μL of well solution (6% ethylene glycol, 8% PEG 8000, 100 mM HEPES pH 7.6) and equilibrated by vapor diffusion against 400 μL of well solution. The harvested crystals were soaked in cryoprotectant solution (well solution supplemented with 20% glycerol) before flash cooling in liquid nitrogen. The oscillation data collection was performed at the National Synchrotron Light Source II beamline 17-ID-1 (AMX) equipped with an Eiger 9M detector at 100 K, a phi of 0.1°, a total of wedge of 180°, and a wavelength of 0.920126 Å.

The diffraction images were reduced using XDS [45]. The R_free_ set was generated from 5% of the reflections in thin resolution shells using Phenix reflection file editor. In the case of Cfl2 and wild-type Cfl1, initial phases were obtained by molecular replacement with a Cif monomer (PDB ID 3KD2, chain A) as a search model using Phaser. In the case of Cfl1-D123S, a monomer of wild-type Cfl1 structure was used as the search model. Iterative automatic (with torsion-angle NCS restraints) and manual model refinement was performed using phenix.refine and Coot [46,47].

### Rosetta Analysis

All calculations were performed using RosettaScripts [23] with Rosetta build version 2020.20.post.dev+53.master.64d08bbfc5f. Briefly, protein models determined by X-ray crystallography were prepared using the FastRelax [24] algorithm with the Rosetta-ICO (a.k.a. beta_nov16) energy function [26] using heavy atom coordinate constraints (see supplementary material for the pre-relax xml protocol); ten decoys were generated and the best scoring model was used for interface analysis. For each protein-protein interface, the shape complementarity [25], change in solvent-accessible surface area (total, polar, and nonpolar area), and binding energy (ΔΔG) were calculated (see supplementary material for the interface analysis xml protocol).

We approximated residue burial using the sidechain neighbor counts method [48]. Briefly, a cone is placed on the c_α_-c_β_ vector for each residue under consideration, and any residue within the defined cone is counted as a neighbor. This metric was calculated through the RosettaScripts interface using the SidechainNeighborCountMetric, which was created for this work.

### Normal Mode Analysis

The Bio3D package as implemented in R was used to carry out NMA using the c-alpha force-field and the nma() command [49,50,51]. The deformation energies displayed in Figure 9 were calculated using the command deformations.nma() [52] and summed over the first three modes for each protein. In the case of the “*ex circulum*” Cfl1 and Cfl2 dimers, the analysis was performed on the dimer subunit that was extracted from its respective oligomer.

### Evolutionary Conservation Analysis

The conservation analysis was performed by the ConSurf online server using the UniRef90 database [27,28]. The multiple sequence alignment used for calculating conservation scores was automatically generated by ConSurf and had to include at least 100 “homologous” sequences to ensure reliability. In the case of aCif, the minimum threshold for percent sequence identity was lowered from 35% to 30% in order to obtain more than 100 “homologous” sequences. The normalized residue conservation scores were binned by ConSurf and colored according to a 9-shade purple-white-green scale.

## Accession Numbers

The coordinates and structure factors for wild-type Cfl1, Cfl1-D123S, and Cfl2 structures are available in the Protein Data Bank under PDB ID: 7JQX, 7JQY, and 7JQZ, respectively.

## Acknowledgements

We would like to thank Dr. Sherry Kuchma for helpful advice on RT-qPCR experiments, and Kelsie Leary for training with CD spectroscopy. We thank Drs. Louisa Howard, Maxime Guinel, and Charles Daghlian for electron microscopy training and assistance. We would also like to thank Drs. Vivian Stojanoff, Jean Jakoncic, Martin Fuchs, Babak Andi, and Wuxian Shi at the FMX, AMX, and SSRL beamlines for advice and support with collection of crystal diffraction data. We are grateful to the members of the Madden lab for helpful discussions and suggestions.

This work was supported by the National Institutes of Health grants R01-AI091699, R01-GM113240, R37-AI83256-06, P20-GM113132, P30-DK117469, and the Cystic Fibrosis Foundation grant STANTO19R0.

