## Supplemental Materials for "Biochemical and Structural Characterization of Two Cif-Like Epoxide Hydrolases from *Burkholderia cenocepacia*"

### Supplemental Materials for Taher *et al.*

The following supplemental materials are included:

- **Table S1** Preliminary screen for Cfl1 and Cfl2 hydrolysis activity against a panel of epoxides.
- **Table S2** Mean ConSurf conservation scores of various interfaces.
- **FIG S1** Comparison of Cfl1 from *B. cenocepacia* strains HI2424 and J2315.
- **FIG S2** Electron microscopy of negatively stained single particles of Cfl2.
- **FIG S3** Quaternary structure differences between Cfl1 and Cfl2.
- **FIG S4** Alignment of the Cfl1-D123S monomer to the Cfl1-WT monomer.
- **Supplemental Movie 1** Comparison of the first non-trivial normal mode of Cif-like proteins.

**Table S1** Preliminary screen for Cfl1 and Cfl2 hydrolysis activity against a panel of epoxides.

| | $\Delta A_{490}$ | | |
|---|---|---|---|
| **xenobiotic substrates** | **Cfl1** | **Cfl2** | **aCif** |
| *S*-styrene oxide | -0.086 | 0.038 | -0.200 |
| *R*-styrene oxide | -0.239 | 0.041 | |
| 1,2-epoxyoctane | -0.037 | -0.037 | |
| 1,2-epoxyhexane | -0.022 | -0.038 | |
| cyclohexene oxide | -0.029 | -0.044 | |
| epibromohydrin | -0.032 | -0.035 | |
| proypylene oxide | -0.016 | -0.036 | |
| glycidol | na | -0.059 | |
| cis-stilbene oxide | na | -0.078 | |
| trans-stilbene oxide | na | 0.006 | |
| 1,2-epoxy-4-vinylcyclohexane | na | -0.041 | |
| allyl glycidyl ether | na | -0.036 | |
| glycidyl methacrylate | na | -0.026 | |
| (+)-limonene oxide | na | -0.045 | |
| **biological substrates** | | | |
| 11(12)-EET | na | -0.039 | |
| 8(9)-EET | na | -0.039 | |
| 16(17)-EpDPA | na | -0.048 | |
| 14(15)-EpETE | na | -0.029 | |
| 5(6)-EET | -0.001 | -0.034 | |
| 14(15)-EET | 0.004 | -0.032 | |
| 17(18)-EpETE | 0.016 | -0.026 | |

The adrenochrome assay was performed as described (see *Materials and Methods* section). $\Delta A_{490}$ values represent the difference in colorimetric readouts between the experimental reactions and the no-substrate reactions, or the D123S catalytic mutant reaction in the case of Cfl1, vs. biological substrates (n=1). Substrate hydrolysis should result in negative values.

**Table S2** Mean ConSurf conservation scores of the non-interface surface residues and of the residues in each monomer:monomer interface of Cfl1 and Cfl2.

| | **residue location** | **<ConSurf score>** |
|---|---|---|
| **Cfl1** | non-interface | 0.47 |
| | 1:2 | 0.06 |
| | 1:3 | 0.28 |
| | 1:4 | 0.13 |
| | 1:7 | 0.44 |
| **Cfl2** | non-interface | 0.42 |
| | 1:2 | 0.25 |
| | 1:3 | 0.21 |
| | 1:4 | 0.07 |
| | 1:9 | 0.76 |

Higher ConSurf scores indicate more variability.

# A

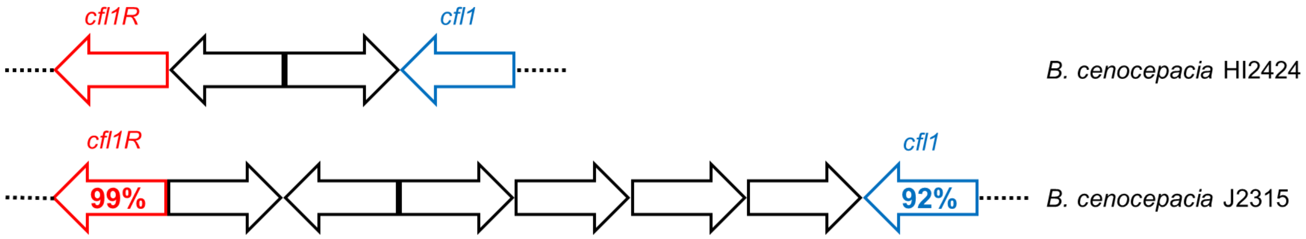


# B

```
Cfl1(H)     MQN---EPSMSGMPAPGLPAGFDRRFSRRYAQVDDVRLHYVTGGPDDGEL      47
            |||   | || |||||||||||:|||||||||:||||||||||||||||:
Cfl1(J)     MQNERSEQSMPGMPAPGLPAGFERRFSRRYAQLDDVRLHYVTGGPDDGEM      50

Cfl1(H)     VVLLHGWPQTWYTWRHVMPVLAQEGYRVVAVDYRGAGESDKPLGGYDKAS      97
            ||||||||||||||||||| ||::||||||||||||||||||||||||||
Cfl1(J)     VVLLHGWPQTWYTWRHVMPALAEDGYRVVAVDYRGAGESDKPLGGYDKAS     100

Cfl1(H)     MAGDIRALVRQLGATRIHLVGRDIGVMVAYAYAAQRPAEIVKLAMLDVPV     147
            ||||||||| ||||||||||||||||||||||||| | ||||||||||||
Cfl1(J)     MAGDIRALVHQLGATRIHLVGRDIGVMVAYAYAAQWPTEIVKLAMLDVPV     150

Cfl1(H)     PGTRIWDEAKARADPQIWHFGLHQQRDIAELLIAGKEHAYILDFYKKRAH     197
            ||||||||||| |||||||||||||||||||:|||||| ||||||||| |
Cfl1(J)     PGTRIWDEAKASADPQIWHFGLHQQRDIAEMLIAGKERAYILDFYKKRTH     200

Cfl1(H)     VALSNDDIAVYADAYAAPGALRAGFELYRAFPQDETQFKAFMKHKLPMPV     247
            |||||||||||||||||||||||||||||||||||||:||||||||||||
Cfl1(J)     VALSNDDIAVYADAYAAPGALRAGFELYRAFPQDETRFKAFMKHKLPMPV     250

Cfl1(H)     LALAGDKSNGAKEFDMAKELALDVRGAVAPNTGHWLPDENPAFLTRQLLD     297
            ||||||||||||| |||:||||||||||||||||||||||||||||||||
Cfl1(J)     LALAGDKSNGAKELDMARELALDVRGAVAPNTGHWLPDENPAFLTRQLLD     300

Cfl1(H)     FFREPAPNR     306
            |||| |   |
Cfl1(J)     FFREAASGR     309
```

**Figure S1.** Comparison of Cfl1 from *B. cenocepacia* strains HI2424 and J2315. (A) The genomic regions that encompass *cfl1-cfl1R* genes are shown. The percent sequence identities of Cfl1 (blue) and Cfl1R (red) homologs in strains HI2424 and J2315 calculated using EMBOSS Needle are indicated on each respective gene. (B) Pair-wise amino-acid sequence alignment of $Cfl1_{(H)}$ and $Cfl1_{(J)}$ calculated using EMBOSS Needle is shown. The subscripts (H) and (J) refer to *B. cenocepacia* strains HI2424 and J2315, respectively. Vertical lines and colons indicate sequence identity and similarity, respectively. The canonical epoxide hydrolase H-G-X-P motif and catalytic residues are highlighted in bold blue and bold red, respectively.

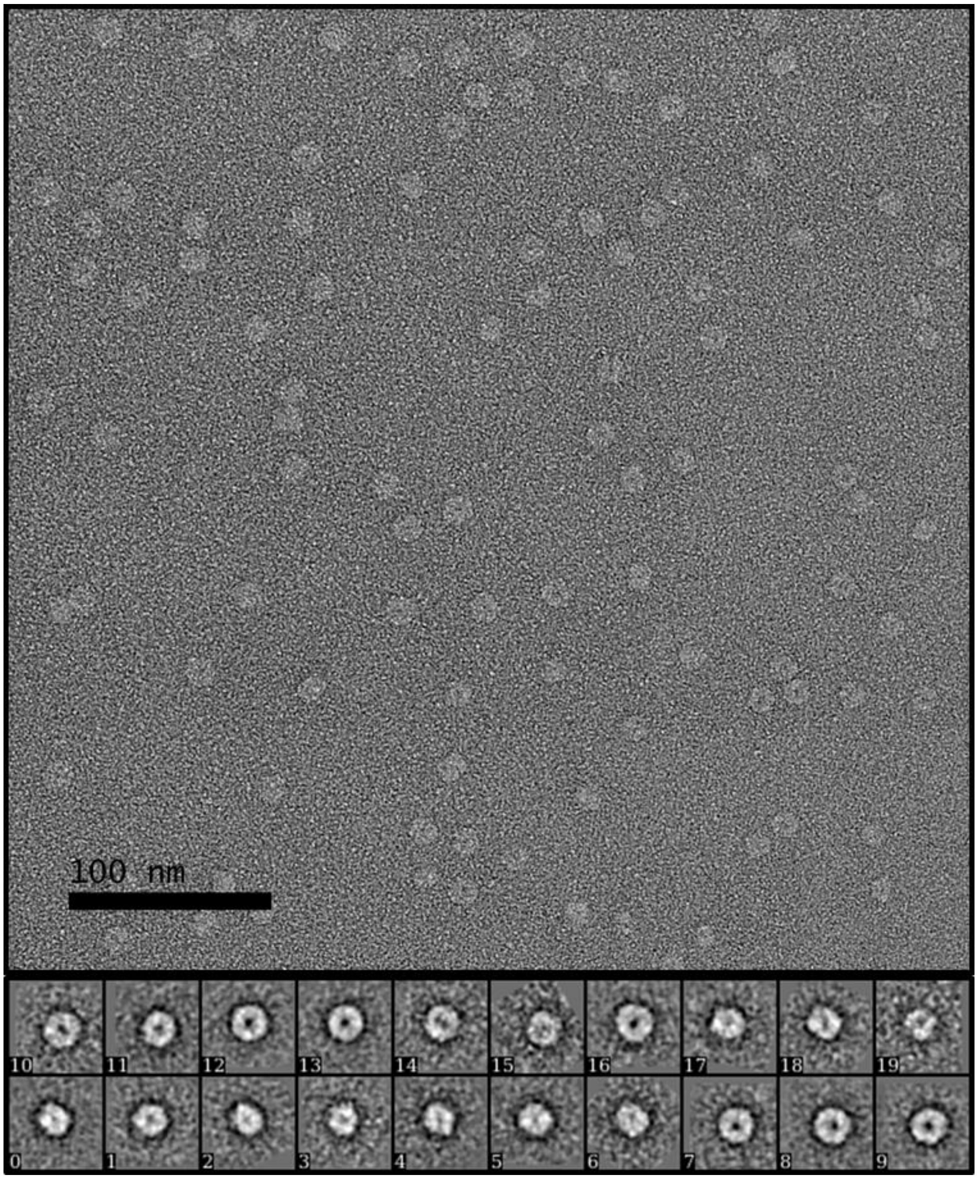


**Figure S2.** Electron microscopy of negatively stained single particles of Cfl2. Top: a representative raw electron micrograph of Cfl2 particles negatively stained with uranyl formate. Bottom: twenty class-average profiles of Cfl2 particles. For experimental details, see *Materials and Methods* section.

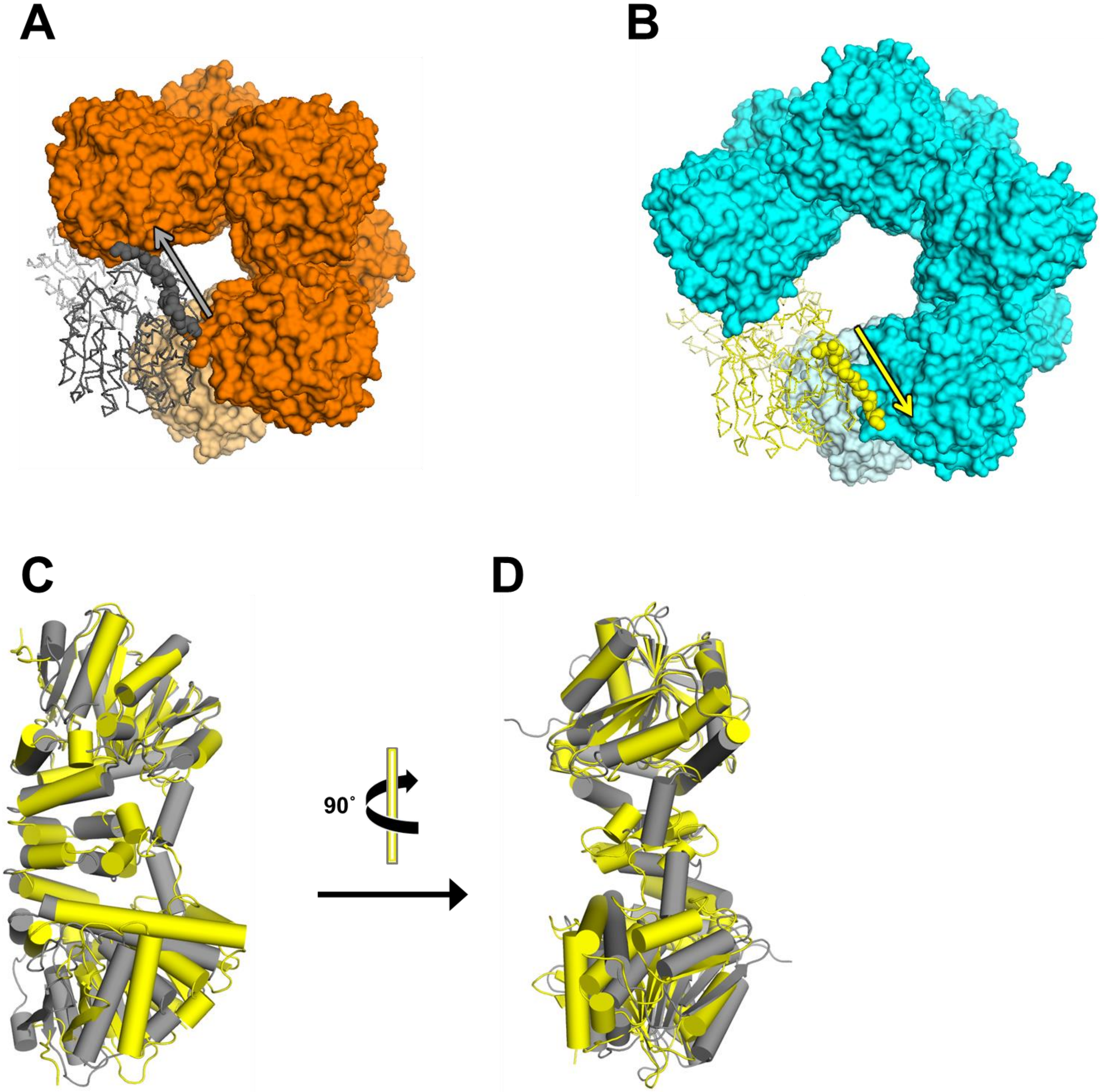


**Figure S3.** Quaternary structure differences between Cfl1 and Cfl2. (A) Top view of the Cfl1 oligomer. Six subunits are shown in surface representation and colored orange, and two subunits are shown as ribbons and colored grey. The N-terminus of one of the subunits is shown as spheres. An arrow indicates the direction the N-terminus extends from one subunit to the other. (B) Top view of the Cfl2 oligomer. Eight subunits are shown in surface representation and colored cyan, and two subunits are shown as ribbons and colored yellow. The N-terminus of one of the subunits is shown as spheres. An arrow indicates the direction the N-terminus extends from one subunit to the other. (C) Side view of the Cfl1 (grey) and Cfl2 (yelow) dimers aligned using the top subunit and shown as cartoon with cylindrical $\alpha$-helices. (D) Same as (C) with a 90˚ rotation about the long axis of the dimers as indicated.

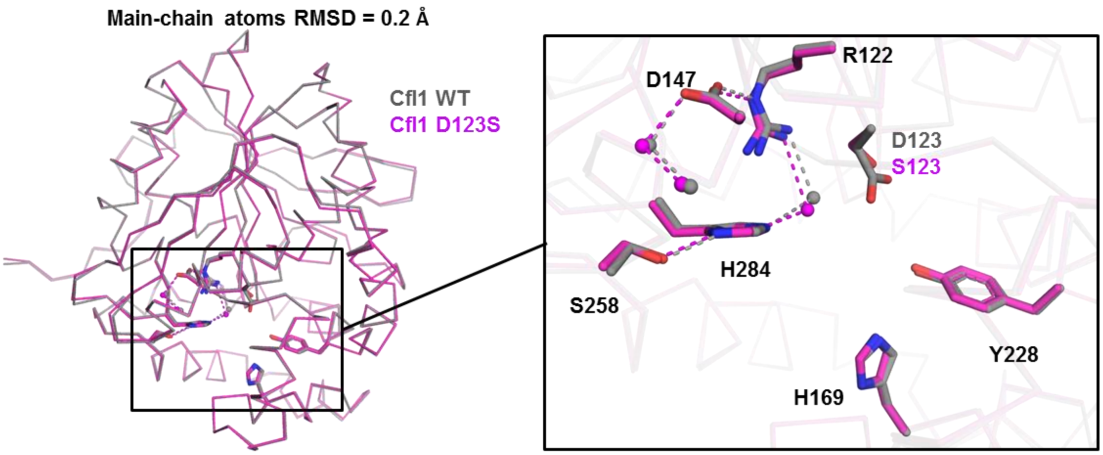


**Figure S4.** Alignment of the Cfl1-D123S monomer to the Cfl1-WT monomer. The main-chain atoms of the Cfl1-WT (grey) and Cfl1-D123S (magenta) monomers were aligned using PyMol. The inset shows a closer view of the presumed catalytic residues (sticks) and waters (spheres). Electrostatic interactions are depicted as dashed lines.

**Supplemental Movie 1** Comparison of the first non-trivial normal mode of Cif-like proteins, including Cif, aCif, and the extracted dimers of Cfl1 and Cfl2. From left to right: Cif, aCif, Cfl1, and Cfl2. The trajectories of mode 7 as calculated by Bio3D are shown for each protein. In the case of Cfl1 and Cfl2, the calculation was performed on the dimer subunit after it was extracted from its respective oligomer (See Materials and Methods for details).
